# Predicting kinase inhibitors using bioactivity matrix derived informer sets

**DOI:** 10.1101/532762

**Authors:** Huikun Zhang, Spencer S. Ericksen, Ching-pei Lee, Gene E. Ananiev, Nathan Wlodarchak, Julie C. Mitchell, Anthony Gitter, Stephen J. Wright, F. Michael Hoffmann, Scott A. Wildman, Michael A. Newton

**Affiliations:** Department of Statistics, University of Wisconsin-Madison, Madison, WI, USA; Small Molecule Screening Facility, Drug Development Core, UW-Carbone Cancer Center, School of Medicine and Public Health, University of Wisconsin-Madison, Madison, WI, USA; Department of Computer Sciences, University of Wisconsin-Madison, Madison, WI, USA; Department of Medicine, University of Wisconsin-Madison, Madison, WI, USA; Biosciences Division, Oak Ridge National Laboratory, Oak Ridge, TN, USA; Department of Biostatistics and Medical Informatics, University of Wisconsin-Madison, Madison, WI, USA; Morgridge Institute for Research, Madison, WI, USA; Department of Oncology, McArdle Laboratory for Cancer Research, University of Wisconsin-Madison, Madison, WI, USA

## Abstract

Prediction of compounds that are active against a desired biological target is a common step in drug discovery efforts. Virtual screening methods seek some active-enriched fraction of a library for experimental testing. Where data are too scarce to train supervised learning models for compound prioritization, initial screening must provide the necessary data. Commonly, such an initial library is selected on the basis of chemical diversity by some pseudo-random process (for example, the first few plates of a larger library) or by selecting an entire smaller library. These approaches may not produce a sufficient number or diversity of actives. An alternative approach is to select an informer set of screening compounds on the basis of chemogenomic information from previous testing of compounds against a large number of targets.

We compare different ways of using chemogenomic data to choose a small informer set of compounds based on previously measured bioactivity data. We develop this Informer-Based-Ranking (IBR) approach using the Published Kinase Inhibitor Sets (PKIS) as the chemogenomic data to select the informer sets. We test the informer compounds on a target that is not part of the chemogenomic data, then predict the activity of the remaining compounds based on the experimental informer data and the chemogenomic data. Through new chemical screening experiments, we demonstrate the utility of IBR strategies in a prospective test on two kinase targets not included in the PKIS. Using limited training data in both retrospective and prospective tests, bioactivity fingerprints based on chemogenomic data outperform chemical fingerprints in predicting active compounds in both standard virtual screening metrics and accurate identification of hits from novel chemical classes.

**Author Summary:** In the early stages of drug discovery efforts, computational models are used to predict activity and prioritize compounds for experimental testing. New targets commonly lack the data necessary to build effective models, and the screening needed to generate that experimental data can be costly. We seek to improve the efficiency of the initial screening phase, and of the process of prioritizing compounds for subsequent screening.

We choose a small *informer set* of compounds based on publicly available prior screening data on distinct (though related) targets. We then use experimental data on these informer compounds to predict the activity of other compounds in the set against the target of interest. Computational and statistical tools are needed to identify informer compounds and to prioritize other compounds for subsequent phases of screening. Using limited training data, we find that selection of informer compounds on the basis of bioactivity data from previous screening efforts is superior to the traditional approach of selection of a chemically diverse subset of compounds. We demonstrate the success of this approach in retrospective tests on the Published Kinase Inhibitor Sets (PKIS) chemogenomic data and in prospective experimental screens against two additional non-human kinase targets.

## Introduction

Early-stage drug discovery involves a search for pharmacologically active compounds (hits) that produce a desired response in an assay on a protein function or disease-related phenotype. The active compounds serve as starting points for further structural optimization, with the ultimate goal of developing therapeutic agents. Virtual screening (VS) can be an effective strategy for prioritizing compounds that can lower high-throughput screening costs by reducing the experimental search to smaller, active-enriched compound subsets localized to the most promising regions of the chemical space. This process can be cheaper and more effective than exhaustive, unguided testing of entire compound libraries [1]. VS also may allow us to evaluate much larger physical or virtual compound libraries. As on-demand synthetic capabilities expand, a VS-guided approach might obviate costs associated with purchasing and on-site storage/maintenance of large general libraries in favor of growing smaller, project-focused compound sets [2].

The choice of which VS methodology to deploy depends on the types of information available at the start of this effort [3]. Structure-based VS methods (such as docking) require specific, structurally-characterized biomolecular targets, but these target structures might only be approximated by homology models [4], or might not be available at all. Phenotypic endpoints like cell death or tumor shrinkage are not amenable to structure-based approaches because specific structural sites of action may not be known. Furthermore, structure-based VS performance varies substantially across targets, where failures are difficult to predict [4,5]. Ligand-based VS approaches can provide more consistent levels of enrichment and are independent from any target structure, but they depend strongly on the quality and abundance of training data in the form of measured compound activities on the target of interest [6]. Such approaches, especially those using topological features for compound representations (such as graph-based fingerprints), may also suffer from high prediction uncertainty when presented with compounds whose chemotypes/scaffolds are outside the scope of the training set [6,7]. The key issue, however, is that training data are usually scarce in early stages of the screening process, making it difficult to generate a predictive model.

For some well-studied target classes (for example, kinases or GPCRs), rich chemogenomic data are available in the form of compound activity profiles across many members of a target class. These data can be structured as a targets-by-compounds matrix of functional interactions, which we term the *bioactivity matrix*. Though sometimes sparse, incomplete, or limited in compound and target coverage, such matrices hold valuable information that can be leveraged for activity predictions on new targets or compounds. We focus on machine learning and statistical strategies that leverage this information to predict the bioactivity profile of a previously uncharacterized member of the target class.

Initial prediction of compound activities may be done by comparing compound features, usually combinations of chemical properties or chemical fingerprints indicating presence/absence of various substructures [6, 7]. Alternative compound fingerprints have been developed on the basis of prior chemogenomic data [8–13]. In these cases, the bioactivity profile of a compound across a series of assays is used as a fingerprint, referred to as a “High Throughput Screening FingerPrint” (HTSFP), based either on continuous bioactivity values or on a binary quantity representing activity/inactivity. HTSFPs enable a useful expression of compound relationships through distances derived among standardized bioactivity profiles in much the same manner as chemical fingerprints.

A key limitation of bioactivity-based inferences is the requirement that some partial profile must be provided for the new target by filling in some compound activities into the new target’s row of the bioactivity matrix. Inference techniques can then be used to fill in the remaining elements in the row. Each element corresponds to an experimental activity measurement for a compound on the new target. So to keep the cost of VS reasonable, the number of experimental measurements required to make accurate predictions of activities on the remaining compounds should be relatively small — a small fraction of the total number of compounds to be prioritized.

Motivated by the need for batch selection strategies to enable effective iterative screening efforts, there has been significant recent effort in developing compound prioritization models from minimal data [14–18]. These methods prioritize additional compounds for testing based on an initial increment of screening data, where the selection of the initial informer subset of compounds to be screened is often random or pseudo-random, or based on chemical diversity. A recent effort by Paricharak et al. [18] uses an active learning process to select an informer set from the most active and least active compounds across a series of PubChem assays. Their work removes specific assay labels from the chemogenomic data to create a balanced data set, and selects compounds on the basis of uncertainty from previous predictive models. However, their optimal informer set is too large to be useful as an initial screening set in most HTS settings.

The focus in these prior studies was iterative model-guided or heuristic-guided batch selection of compounds for multiple phases of screening, rather than optimal selection of the informer set of compounds to be labelled for the new target. Our emphasis in this paper is on selection of the *informer set* — the initial (0^th^ iteration) set of compounds to be assayed in a multi-phase scheme. We refer to approaches based on informer sets as *Informer-Based Ranking* (IBR) methods. Our approaches are analogous to earlier chemometric experimental design approaches like chemical cluster sampling [19], but leverage chemogenomic data instead. The proposed IBR methods each involve two steps; see Figure 1. In the first step, we select an informer set of compounds to evaluate experimentally for bioactivity on the new target. Importantly, this selection is guided by the bioactivity matrix. The second step involves prioritization of the compounds outside the informer set, according to their bioactivity against the new target. This prioritization may make use of both the bioactivity data on other targets as well as the new data obtained on the informer compounds on the new target. We describe algorithms both for the selection of the informer compounds and for the prioritization of other compounds after screening data are obtained for the informer compounds.

**Fig 1.**
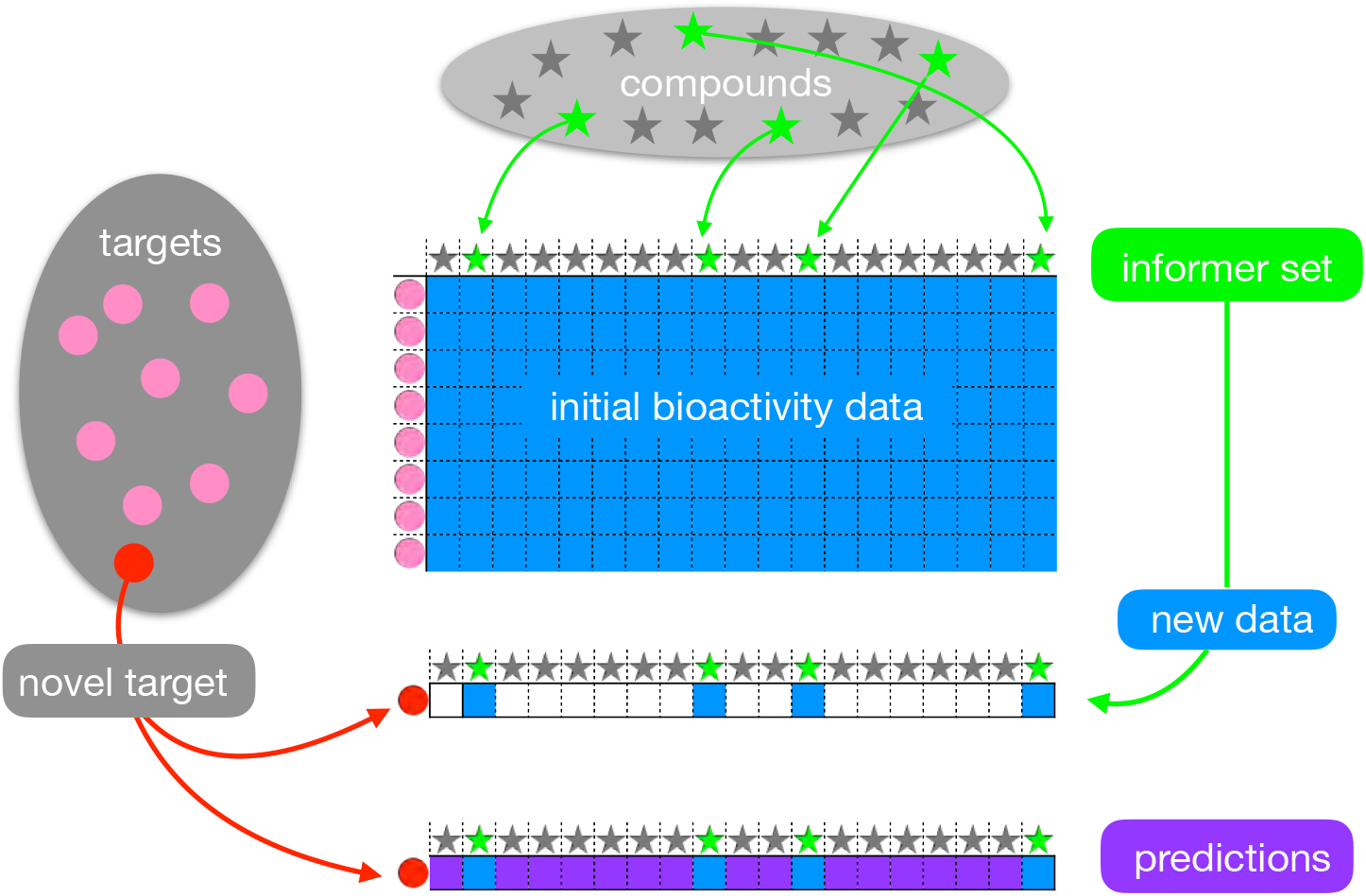
IBR (Informer-Based Ranking) for compound prioritization on a novel target. From a complete bioactivity data matrix (blue grid), a subset of informer compounds (green stars) are identified from the broader set of compounds (stars) that have been tested against a large set of targets (pink circles). A previously uncharacterized target (red circle) is assayed with just the informer compounds, and the new bioactivity data are used to reveal the new target’s relationship to other targets. The combined data enable activity predictions (purple) on the remaining, non-informer compounds.

We propose three novel IBR strategies: Regression Selection (RS), Coding Selection (CS), and Adaptive Selection (AS). Each strategy consists of (i) an informer set selection method that chooses a small number of compounds to be tested, based on characteristics of the bioactivity matrix, and (ii) a compound ranking method that leverages returned informer data to predict which of the untested compounds is active against a new target not represented in the currrent bioactivity matrix. Underlying all three strategies is the premise that targets may be naturally organized according to patterns in their bioactivity profiles across compounds. This organization leads to a clustering of targets as well as to the identification of informer compounds that are predictive of the cluster identity of a novel target. The strategies leverage advances in optimization and statistical analysis, and they differ in how patterns are recognized and computations are deployed.

We apply the proposed IBR strategies in the context of two public human kinase chemogenomics matrices: PKIS1 [20] and PKIS2 [21]. We deploy the strategies prospectively, by prioritizing PKIS1 and PKIS2 compounds for activity against two distinct kinase targets of potential therapeutic importance: *Mycobacterium tuberculosis* PknB and Epstein-Barr BGLF4. We also apply the strategies retrospectively, in a cross-validation study of each chemogenomics matrix, leaving out one target at a time and prioritizing compound activity against the left-out target.

The performance of each new IBR strategy was assessed prospectively by inspection of the successful activity predictions, and retrospectively using common VS metrics, including: Area Under the Receiver-Operator Characteristic Curve (ROCAUC) and enrichment factor (EF). We also assessed each strategy’s ability to retrieve structural diversity among active compounds by computing the fraction of active scaffolds identified in the top of the ranking. Further, we compared the proposed IBR strategies to several simpler IBR approaches, including ones that use compound structure information and similarity-based expansion, and ones that use marginal features of the bioactivity matrix.

## Results

### IBR strategies apply in the low-data regime

We describe IBR strategies that require experimental testing of some new target of interest on a small fraction of the compound library — the informer subset — with a view to effectively prioritizing the remaining compounds for subsequent testing for activity with the target. The complete IBR strategy thus has two parts: a scheme to identify the informer subset and a scheme to prioritize the remaining compounds after assay data have been obtained for the informer compounds. Initially, we may have no assay data on the new target, though we typically have some such chemogenomic data on related targets that populate a related sector of chemical space, in some sense. Ideally, a successful IBR strategy might be applied in target-agnostic drug development settings (for example, phenotypic targets or incompletely featurized targets), so we intentionally exclude from each IBR strategy target-specific features, such as protein sequences or structural information.

We described three novel IBR strategies that utilize statistical patterns in the bioactivity matrix that is available prior to informer-set assay testing. Regression Selection (RS), Coding Selection (CS), and Adaptive Selection (AS) all treat the target space as being partitioned into clusters of targets so that, within each cluster, there is some relevant similarity of the bioactivity profiles of the targets across the space of tested compounds. These three strategies also posit that a small number of compounds (the informer subset) have bioactivity profiles that are predictive of the cluster label appropriate to any target, including the novel target of interest. RS, CS, and AS differ in how they evaluate clusterings and potential informer subsets. For example, RS and AS involve kmeans clustering of targets followed by regularized multinomial regression to learn the relationship between compounds and cluster labels, but they differ in how the regression is regularized and how the informer compounds are identified. In contrast, CS forms a single objective function that simultaneously scores clustering strategies and potential informer compounds.

Computationally simpler baseline IBR strategies are useful to consider, as they may approximate practical experimental design scenarios. <u>B</u>aseline <u>C</u>hemometric strategies (BC_s_, BC_l_, and BC_w_) use chemical features for both informer selection and non-informer ranking. Three different chemometric ranking strategies are used for the non-informer ranking, as denoted by subscripts s, l, and w (described in detail in the Methods). Here, clustering is applied on the compound space using the known chemical structure (fingerprints) of the compounds (not used in RS, CS, or AS) in order to identify informer compounds. Then, prioritization of the non-informers makes use of various ways of ranking the chemical distance between bioactive informers and non-informers. Alternatively, a <u>B</u>aseline *<u>F</u>requent-hitters* strategy simply takes as informer compounds those that show the highest rate of activity within the initial target set (BF_s_, BF_l_, BF_w_). Prioritization of non-informers uses chemical distance, as in the chemometric methods. To simplify, we only report baseline results for each of our top chemometric and frequent-hitters baseline strategies (BC_w_ and BF_w_). Outcomes for the full set of baselines are available in Supplemental Information.

Performance of the IBR strategies was evaluated using two virtual screening metrics that reflect successful prioritization of active compounds ROCAUC and <u>N</u>ormalized <u>E</u>nrichment Factor in top <u>10</u>% of ranking (NEF10)). An additional metric <u>F</u>raction of <u>A</u>ctive <u>S</u>caffolds <u>R</u>etrieved (FASR10) assesses the diversity of the active chemical structures that were prioritized in the top 10% of the ranking.

### Prospective tests of IBR strategies on novel kinase targets

We applied the IBR strategies on two novel kinase targets outside of the PKIS1 and PKIS2 target sets. These targets are phylogenetically distant from most of the human protein kinases in the PKIS data sets, with relatively low sequence identity to the nearest neighbors in the PKIS1/2 sets in comparison to kinase domain sequences (Supplementary Figure S1). For *Mycobacterium tuberculosis* kinase PknB (UniProt ID: P9WI81), the nearest neighbors were the human serine/threonine kinases MARK2 (16.1% sequence identity) in PKIS1 and BRSK1 (16.1%) in PKIS2 (UniProt IDs: Q7KZI7 and Q8TDC3, respectively). For Epstein-Barr virus kinase BGLF4 (UniProt ID: I1YP37), the most similar sequences were from human protein tyrosine kinase (PTK2 or FAK2) (13.8%) in PKIS1 and human serine/threonine-protein kinase (LRRK2) (14.2%) in PKIS2 (UniProt IDs: Q14289.2 and Q5S007).

To prioritize which PKIS compounds might be active on PknB and BGLF4, each IBR strategy selected 16 informer compounds from PKIS1 and 16 informer compounds from PKIS2 that were assayed on PknB or BGLF4 using recombinant kinase proteins and the ADP Glo reagent. Informer set selections by each IBR for PKIS1 are shown with their associated experimental bioactivity measurements in Supplementary Table S1. The assay results for the informer compounds selected by each of the IBR strategies were used to rank the remaining non-informers in PKIS1 or PKIS2. To evaluate the performance of the different methods, all of the available PKIS1 and PKIS2 compounds were assayed on PknB or BGLF4. Experimental active/inactive labels were assigned using *μ*+2*σ* percent inhibition (activity) thresholds in PKIS1: PknB=13.4% and BGLF4=20.2%, and PKIS2: PknB=8.7% and BGLF4=12.5%, based on screening results from the PKIS compound sets.

The RS and CS approaches were the only methods that recovered multiple hits and active scaffolds in their top 10% of ranked compounds for both kinase targets and both datasets (Table 1). RS managed to recover actives for PknB even though it did not include any active compounds in its PKIS1 or PKIS2 informer sets (Table S2). The RS method was also the best overall for BGLF4 on PKIS2, and tied as the best method for PknB on PKIS1. CS was the best approach for PknB on PKIS2.

**Table 1.** Retrieval counts by the various methods on new kinase targets (a) PknB and (b) BGLF4 using PKIS1 or PKIS2 matrices. The total number of experimentally determined active compounds and scaffolds is indicated in the *total* column. The values below each of the IBR methods indicate the experimentally determined active compounds that were ranked in the top 10% of predicted active compounds by each method.

| matrix | hits | baselines |  | non-baselines |  |  | total |
| --- | --- | --- | --- | --- | --- | --- | --- |
|  |  | BC<sub>w</sub> | BF<sub>w</sub> | RS | CS | AS |  |
| PKIS1 | compounds | 1 | 7 | 7 | 2 | 3 | 8 |
|  | scaffolds | 1 | 7 | 7 | 2 | 3 | 7 |
| PKIS2 | compounds | 0 | 1 | 2 | 3 | 1 | 7 |
|  | scaffolds | 0 | 1 | 2 | 3 | 1 | 7 |

(b) BGLF4
| matrix | hits | baselines |  | non-baselines |  |  | total |
| --- | --- | --- | --- | --- | --- | --- | --- |
|  |  | BC<sub>w</sub> | BF<sub>w</sub> | RS | CS | AS |  |
| PKIS1 | compounds | 3 | 9 | 3 | 7 | 10 | 11 |
|  | scaffolds | 2 | 6 | 3 | 5 | 7 | 8 |
| PKIS2 | compounds | 1 | 1 | 8 | 3 | 1 | 10 |
|  | scaffolds | 1 | 1 | 7 | 3 | 1 | 8 |

AS and the three BF baseline methods (BF_w_ shown in Table 1) struggled for both targets with the PKIS2 compounds, each identifying only a single hit. However, AS was the best approach for BGLF4 on PKIS1 compounds. Finally, the three purely chemometric baseline approaches (BC) (BC_w_ shown in Table 1) were the worst overall, in many cases failing to recover any hits. The best methods were the same when evaluated with NEF10 or FASR10, but varied slightly for ROCAUC (Table S3).

### Retrospective tests of IBR strategies by cross validation on PKIS1 data matrix

As a further assessment of IBR methods, we conducted retrospective leave-one-target-out (LOTO) analysis for each of the *m* = 224 targets in PKIS1. This involved *m* = 224 separate applications of all the IBR strategies applied to reduced chemogenomics matrices (*m* − 1 rows), again using an informer size of 16 compounds. Each time, the bioactivity profile of the left-out target was predicted in the sense that compounds were prioritized for activity against this one left-out target.

Results from PKIS1 LOTO cross validation are summarized in Table 2. With respect to the ROCAUC metric (Figure 2), the purely bioactivity-based *RS* model provides the best rankings with a median ROCAUC value of 0.92 ± 0.11 (± one standard deviation). *RS* and *AS* methods both had better performance than the top chemocentric and frequent-hitter baseline approaches, BC_w_ (0.67 ± 0.22) and BF_s_ (0.83 ± 0.14). The improvements in ROCAUC of *RS* and *AS* over BC_w_ and BF_s_ were statistically significant (all *p*-values < 1 × 10^−5^). The *CS* method also had statistically better ROCAUC performance than all baseline models except BF_l_ (*p* = 0.053). A complete set of *p*-values from a pairwise comparison of the IBRs is available in Table S5. The hybrid baseline approaches, which use compound bioactivity profiles to select the most broadly active compounds as informers, performed much better than the chemometric approaches that use chemical features for informer selection.

**Fig 2.**
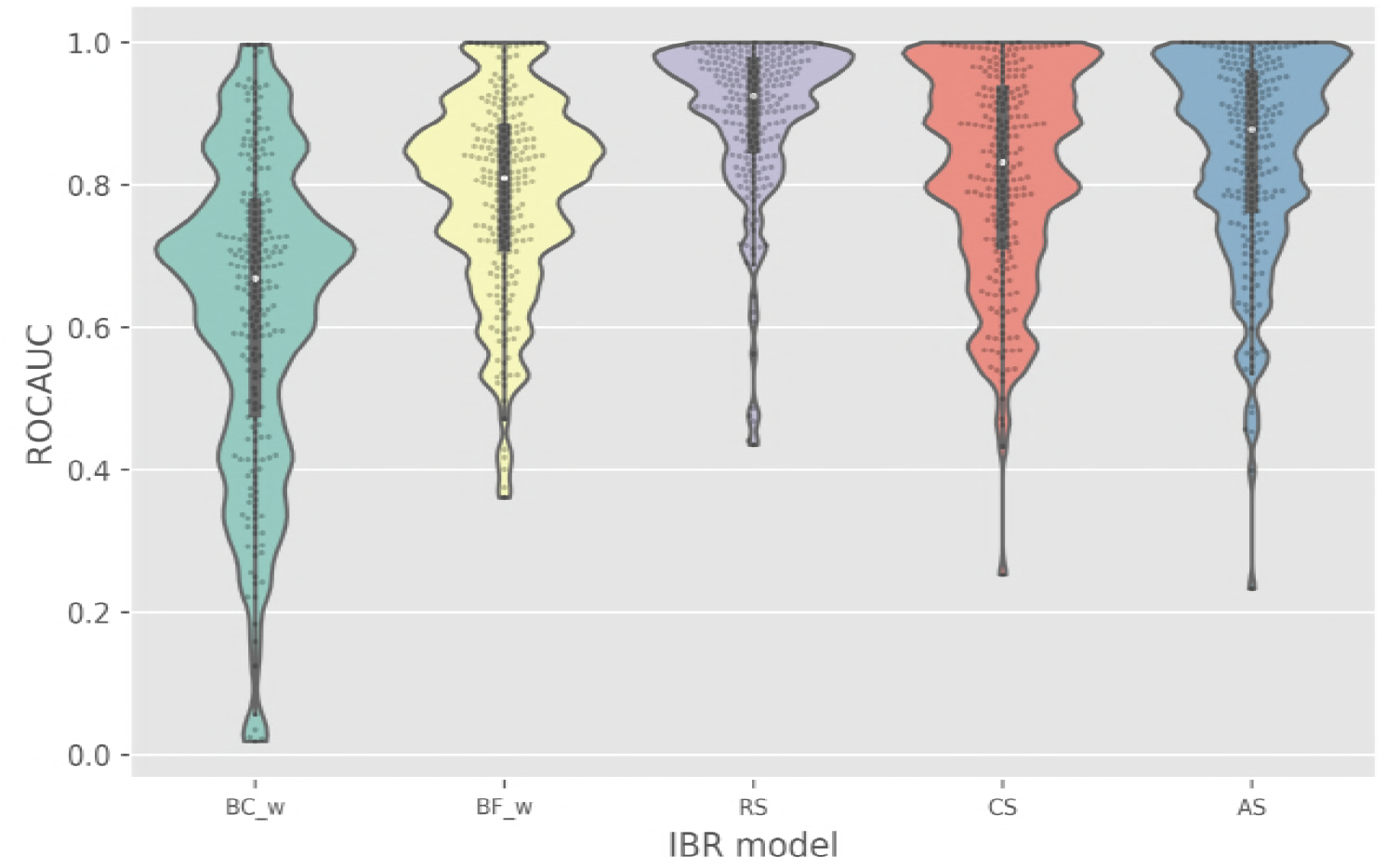
A comparison of models with respect to compound ranking performance as assessed by ROCAUC values. Each model was evaluated on 224 targets through PKIS1 leave-one-target-out validation. ROCAUC of 0.5 indicates a random ranking of compounds on a given target; ROCAUC of 1.0 represents ideal ranking with all active compounds prioritized above the inactives. The individual target evaluations are shown as light grey dots and the median and interquartile range are displayed as a white circle and black bars, respectively.

**Table 2.** (a) ROCAUC, (b) NEF10, and (c) FASR10 in Leave-One-Target-Out Cross Validation on PKIS1. IBR methods were evaluated on 224 PKIS1 targets using standard VS metrics that reflect active retrieval: ROCAUC and NEF10. FASR10 was also evaluated to reflect the chemical diversity of the actives retrieved. All baseline outcomes are shown in Figure S4 along with *p*-values from pairwise comparisons in Table S5. *The only non-baseline IBR that fails to demonstrate statistical improvement (*p* <0.05) over all baselines is CS when using the ROCAUC metric. A Šidák multiple comparison correction was applied using 6 baselines against each non-baseline IBR.

|  | baselines |  | non-baselines |  |  |
| --- | --- | --- | --- | --- | --- |
|  | BC<sub>w</sub> | BF<sub>w</sub> | RS | *CS | AS |
| mean | 0.63 | 0.79 | 0.90 | 0.81 | 0.84 |
| median | 0.67 | 0.81 | 0.93 | 0.83 | 0.88 |
| stdev | 0.21 | 0.13 | 0.11 | 0.14 | 0.14 |

(b) NEF10
|  | baselines |  | non-baselines |  |  |
| --- | --- | --- | --- | --- | --- |
|  | BC<sub>w</sub> | BF<sub>w</sub> | RS | CS | AS |
| mean | 0.62 | 0.74 | 0.80 | 0.79 | 0.82 |
| median | 0.60 | 0.72 | 0.81 | 0.79 | 0.85 |
| stdev | 0.13 | 0.13 | 0.13 | 0.14 | 0.13 |

(c) FASR10
|  | baselines |  | non-baselines |  |  |
| --- | --- | --- | --- | --- | --- |
|  | BC<sub>w</sub> | BF<sub>w</sub> | RS | CS | AS |
| mean | 0.31 | 0.52 | 0.68 | 0.65 | 0.71 |
| median | 0.29 | 0.50 | 0.72 | 0.64 | 0.75 |
| stdev | 0.21 | 0.21 | 0.22 | 0.26 | 0.23 |

We also compared strategies using enrichment factor (EF) as an alternative VS metric that, like ROCAUC, reflects retrieval of active compounds (Figure 3). The maximal EF value that could be achieved on a target, however, depends on the active fraction in the set. To address the variation in the extent of the class imbalance across kinase targets (active fractions ranging from 0.01-0.12 in PKIS1) (Figure S2), we apply the normalized EF metric NEF10. The EF cutoff was also extended from a typical 1% threshold out to 10%, due to the small number of compounds considered (*n* = 366). To simplify comparison with the ROCAUC metric, we scale NEF10 such that a value of 0.5 reflects a random classifier (equivalent to random ranking or no enrichment) and a value of 1.0 represents a perfect classifier, in which the top 10% has been maximally enriched. Over the 224 targets considered in PKIS1, our three bioactivity-based models (RS, CS, and AS) are statistically superior to all of the baseline approaches (all *p* < 1 × 10^−13^). The AS method had the strongest enrichment for active compounds, with median NEF10 of 0.85 ± 0.13. This was better than the top frequent hitters model, BF_l_, which had a median NEF10 of 0.74 ± 0.13 (*p* < 1 × 10^−9^). The enrichment is even better compared to the chemometric models, the best of which is BC_w_, providing a median NEF10 of 0.60 ± 0.13 (*p* < 1 × 10^−9^).

**Fig 3.**
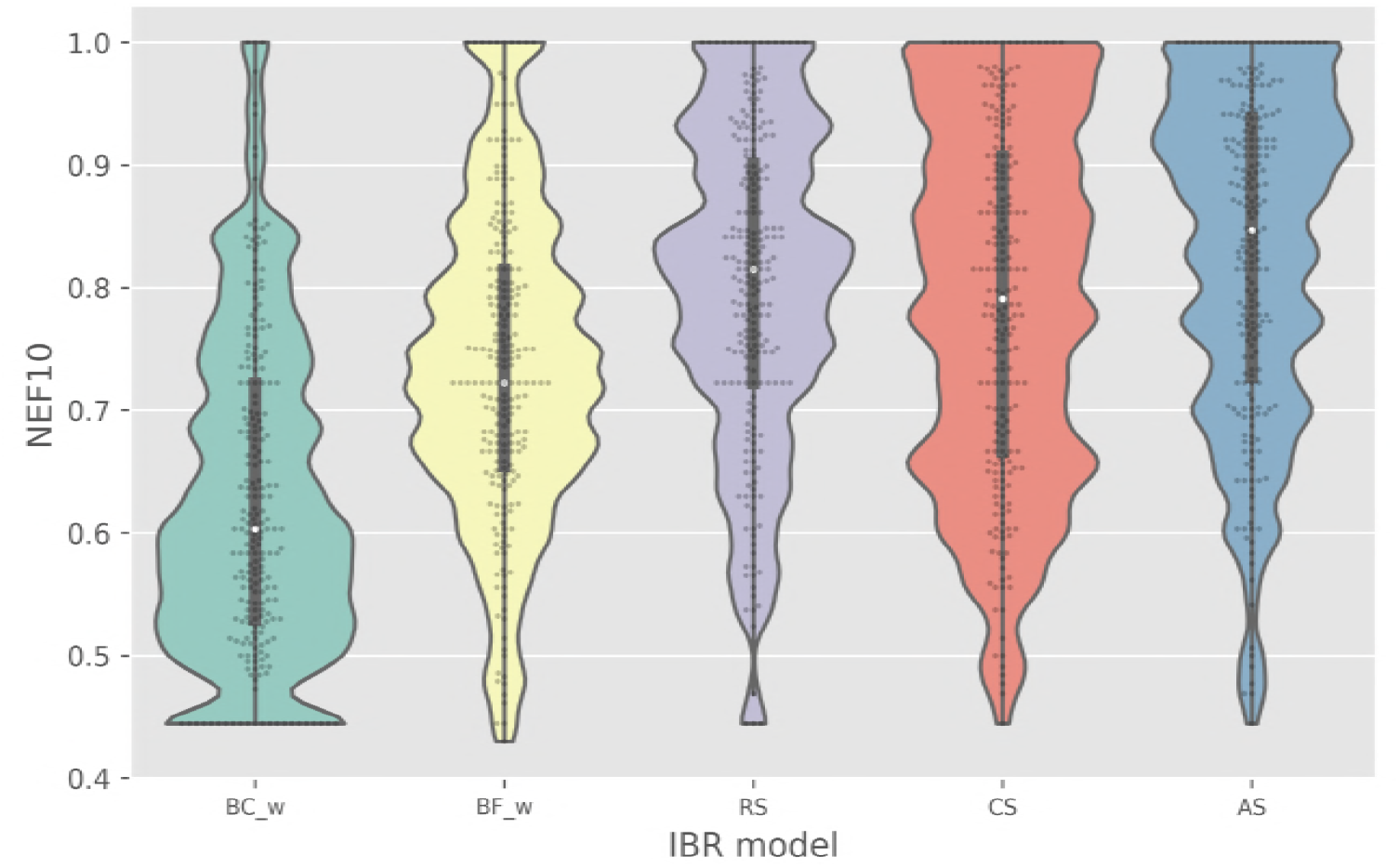
A comparison of models with respect to compound ranking performance as assessed by active enrichment in the top 10% of ranked compounds. Each model was evaluated on 224 targets through PKIS1 leave-one-target-out validation. NEF10 represents the fold-enrichment of actives in top 10% above random that is normalized by dividing by the maximum theoretical fold-enrichment that could be achieved at the 10% threshold for the target of interest.

Another key characteristic of robust virtual screening performance is the recognition of diverse active compound structures, rather than retrieval of only a subset of the active chemotypes. Because of the high rate of failure for hits in follow-up hit-to-lead or optimization efforts, we value methods that can retrieve as many active scaffolds as possible, even at some expense to predictive accuracy reflected by ROCAUC and NEF metrics. Across PKIS1 targets, we assessed the diversity among the known active chemotypes prioritized by each model by monitoring the Fraction of Active Scaffolds Retrieved among the top ranking 10% of compounds (FASR10) (Figure 4). The bioactivity-based IBR methods outperform the top hybrid and chemocentric baseline models, according to this metric. The median FASR10 for the *AS* model 0.75 ± 0.23 exceeded the top hybrid model, BF_l_ (0.53 ± 0.22), and chemocentric model, BC_w_ (0.29 ± 0.21), (with *p* < 1 × 10^−9^ and < 1 × 10^−9^, respectively).

**Fig 4.**
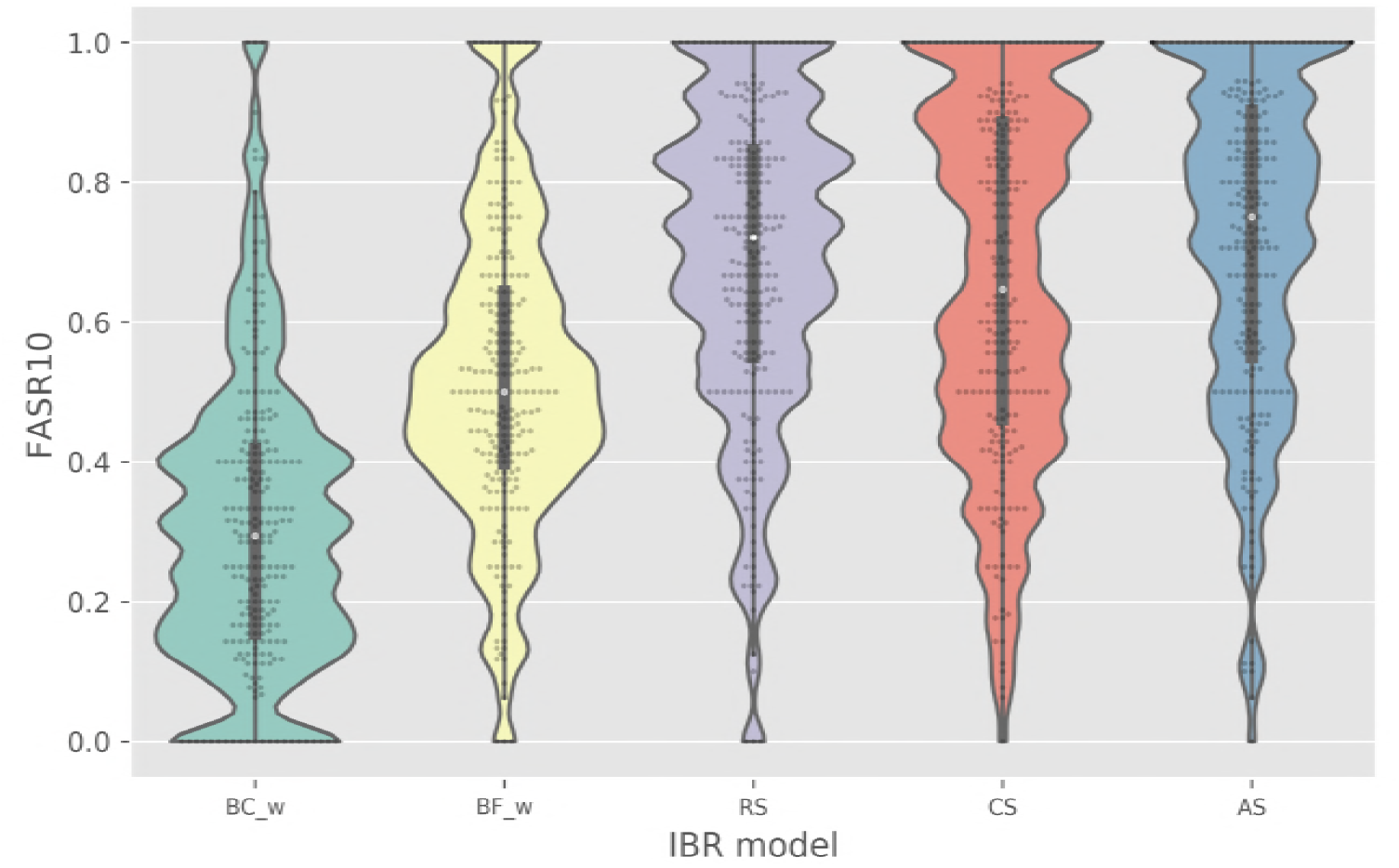
A comparison of models with respect to the structural diversity of the active compounds retrieved. Each model was assessed by FASR10 evaluations on 224 targets through PKIS1 leave-one-target-out validation. The FASR10 metric is the fraction of the total identified active molecule scaffolds, for the target of interest, that were identified in the top 10% of the ranked compounds on that target. Compounds are grouped by their generic (all-carbon skeletons) representations of Bemis-Murcko scaffolds.

### Effect of Informer Set Size

All IBR strategies require choosing the number of elements *n_A_* to include in the informer set. Larger *n_A_* allows more information to be gleaned from intermediate screening data, and therefore improved prioritization of non-informer compounds. Marginal improvements in performance as a function of *n_A_* are expected to diminish as *n_A_* increases, because of redundancies in the information gleaned as more activity data accrues. Larger *n_A_* also leads, of course, to higher assay costs. The experiments reported above used *n_A_* = 16, amounting to about 4% of the compounds in the chemogenomics matrix.

To examine the relationship of informer set size to prioritization performance, we applied IBR strategies on a range of informer set sizes. First, we considered AS, our best performing IBR strategy. Figure S4 shows ROCAUC and NEF10 metrics from the LOTO retrospective analysis of PKIS1 for *n_A_* varying from 9 to 28. Performance did not vary greatly over this range. We also tested a wider range of informer set sizes (*n_A_* = 1 to 48 compounds) on PKIS1 target predictions using LOTO cross validation, and examined ROCAUC and NEF10 using baseline IBR methods BC_w_ (Figure S5) and BF_w_ (Figure S6). Over this range, we observe performance degradation with diminishing informer set sizes. These experiments indicate that our preferred value *n_A_* = 16 strikes a reasonable balance between size and performance for this particular data set.

## Discussion

We set out to establish effective strategies to prioritize compounds for initial testing in iterative high-throughput screens in a drug discovery setting. Our approach is related to the cold-start problem in collaborative filtering (recommender systems) and involves informer-based ranking (IBR) strategies that identify a small subset of highly informative compounds to test in the initial screening round. Data obtained by testing the informers can be used to prioritize compounds for subsequent screening. As a proof of concept, we focused on kinases, so that we could test methods using public kinase chemogenomic data matrices. Among the IBR strategies tested, we found that those that leveraged bioactivity data from matrix targets (RS, CS, and AS) provided better initial sampling than baseline strategies that applied chemometric similarity methods (BC) or hybrid approaches (BF). (The hybrid approaches used a “frequent hitters” heuristic for informer selection, based on matrix activities, and chemometric similarity for ranking.)

We applied our chemogenomic IBR and baseline methods in prospective tests on two microbial kinases: PknB and BGLF4. An initial batch of just 16 informer compounds from each set (roughly 4% of the complete set of compounds) was selected for assays on PknB and BGLF4. The methods were evaluated with regard to hit prioritization and diversity of active scaffolds prioritized, compared to the results of assays of the PKIS1 and PKIS2 compounds. Results from these prospective tests indicated that IBRs using bioactivity data and hybrid baseline IBRs outperformed baseline IBRs that use purely chemometric data.

For a more complete assessment of the IBRs, we performed a retrospective leave-one-target-out validation on the PKIS1 matrix (*m* = 224 targets by *n* = 366 compounds) using a batch selection of 16 informer compounds. We observed statistically better hit prioritization and active scaffold retrieval for the purely bioactivity-based IBRs (RS, CS, and AS) than for any of the baseline methods.

### IBRs and Related Approaches

Chemogenomic assay data have been used through inductive transfer or transfer learning approaches to make successful predictions on compound-target interactions in several contexts [22,23]. Reker et al. [16] and Cichonska et al. [24] placed chemogenomic predictions into 4 classes: (1) filling in missing elements within a relatively complete chemogenomic matrix (bioactivity imputation), (2) predicting interactions for a target on matrix compounds (virtual screening), (3) predicting interactions for a compound on matrix targets (drug re-purposing or off-target effects), and (4) predicting interactions for non-matrix compounds on a non-matrix target (virtual screening). Wasserman et al. [25] showed that simple kernel approaches using nearest proxy targets could be used to rank compounds effectively for a query target (class 2), as long as it was possible to identify proxy targets closely related to the query target. For kinase targets, Cichonska et al. [24] explored a wide range of ligand and target kernels to address class-1 and class-3 problems. For focused target sets (kinases and GPCRs), Janssen et al. [26] recently applied nearest-neighbor approaches to ligand and targets mapped on t-SNE projections to address class-2 and class-3 problems.

The methods we report differ from prior chemogenomic methods for addressing the class-2 problem by involving strategic but limited data acquisition on the query target. Determination of the responses of targets to key informer compounds shifts a relatively difficult class-2 problem into the more tractable class-1 problem of imputation. Unlike chemogenomic kernel-based approaches [24, 25], we did not use target features, focusing instead on target-agnostic strategies for compound ranking that could be used in the future for cell-based or phenotypic assays. Our focus on limited, strategic data acquisition on the target of interest frames the problem in a more practical context akin to compound prioritization in early, low-data stages of an iterative screening effort [14,18,27,28]. Lack of active compound instances can stall implementation of supervised models for compound selections [15]. Our bioactivity-based IBR methods overlap hit expansion methods using chemogenomic data, as applied by groups at Novartis for guiding molecule selection in iterative screening [18,27]. In agreement with their findings, IBR methods that use compound bioactivity profiles, rather than chemical features, provided broader active scaffold retrieval [27]. Previous implementations of HTSFP, however, define compounds by normalized bioactivity vectors from an independent reference assay set, whereas our IBRs use compound bioactivity profiles derived directly from the available chemogenomic matrix. We tested targets only from the same target class, namely, protein kinases. The IBR-based informer sets could be applied in the same way that Paricharak et al. used their Mechanism-of-Action Box (MoABox) of probe compounds for testing in “iteration zero” of their iterative screening procedure [27].

### Practical Implications

The IBR strategies described here could enable iterative screens either on orphan members of a target class or on targets on which very few compounds have been tested. Data returned on each screening iteration would then be used as new training instances to refine the model, potentially in an active-learning framework that also considers relevance of training instances for subsequent compound selection. To promote efficiency of an iterative approach, initial compound batches are often limited in size, with compounds are often being selected at random or to achieve chemical diversity. Initial screens chosen in this way are likely to return few active compounds, thus stalling effective implementation of a supervised activity prediction model. The IBR strategy reported here can be deployed for compound prioritization in early rounds of batch selection; the informer set could be tested to obtain preliminary compound rankings in the low-data phase of iterative screens. Due to class imbalance being skewed towards inactive compounds in drug discovery tasks, IBR methods could enable rapid identification of relatively rare but important active instances necessary for training the activity-prediction model until it can score compounds accurately for prioritization.

Moreover, the bioactivity-based IBR methods exhibited diverse active-scaffold recognition properties, yielding positive training instances with greater structural diversity for supervised compound prioritization models. The FASR10 results indicate that bioactivity-based IBR approaches generalize better over different compound structures than chemometric IBRs, so they should exhibit a greater tendency to scaffold hopping [27,29,30]. In contrast, all of the baseline IBR methods use Morgan fingerprint-derived distances to active informers, thus confining their perspective to those active regions of chemical space identified with the informer set. Different chemotypes, however, can exhibit strong activity on the same target. Plots of PKIS1 compounds projected into their three major principal components of chemical feature space (Morgan fingerprints) frequently show active regions that are non-adjacent (Figure S3). While active compounds tend to cluster in specific regions of chemical space, many targets elicit multiple, sometimes distantly separated regions of active chemical space.

### Future Directions

There are several potential uses for IBRs in drug discovery. This work demonstrates the possibility of effective prediction of activities for new targets within the same target class (kinases) from an extensive chemogenomics data matrix representing many targets within that class. A future direction of research is to quantify the amount of chemogenomic data needed to enable robust prediction within the same target class. It appears that low-rank structure in the chemogenomic matrix used in the IBR methods helps to enable reliable predictions of a target’s compound preferences. Statistical models that faithfully represent variation and dependence in bioactivity data also could be leveraged to guide the development of alternative IBR strategies beyond RS, CS, and AS.

Of greater interest is the development of a more general informer set from a broader collection of chemogenomic data. To investigate the generalizability of the methods, we plan to apply them to a wider range of novel targets (or held-out targets) using an expanded chemogenomic data set with broader target and compound coverage. We do not know how well IBRs will perform on new targets that are unrelated to those within the matrix. We are encouraged by the prospective predictive performance on query kinases (PknB and BGLF4) that are dissimilar from kinases in the chemogenomic data, but note that these targets are still related functionally. More comprehensive data matrices tend to be incomplete, with many missing data values, but they should be useful in testing whether these methods are effective in extended pharmacological spaces. The size of the informer set may well have a dramatic impact on overall performance.

It may be possible to use IBR methods for prioritizing non-matrix compounds on a new target (a class-4 problem). Chemogenomic matrices enable pharmacological mapping of a given new target (query) to matrix targets that exhibit similar bioactivity profiles (proxy targets). Associations between query targets and proxy targets can be made on the basis of full-compound bioactivity profile in the matrix, or potentially just informer assay results. Given that certain proxy targets are likely to be more extensively screened (tested with compounds outside the matrix set), it might be possible to use non-matrix screening data on proxy targets to infer activities for additional compounds and thus prioritize them for testing on some query target.

## Materials and Methods

### PKIS data sets

Most of the IBR strategies developed here leverage chemogenomics data matrices for activity predictions on compounds against selected kinase targets. The matrices were derived from two public human kinase chemogenomics data sets PKIS (PKIS1) [20,31] and PKIS2 [21]. Prior to development and testing of methods, these sets were processed as described below. (Links to our processed PKIS datasets are provided below.)

*PKIS1*. The original PKIS data set (PKIS1) was downloaded from https://www.ebi.ac.uk/chembldb/extra/PKIS/PKIS_screening_data.csv Each row in this data set contains an assay result on a specific compound. Each row lists several identifiers for each compound and the target, assay conditions, and the assay read-out (percent inhibition). For nearly every compound, kinase activity was tested independently at 0.1 *μ*M and 1.0 *μ*M concentrations. For this work, only the inhibition values obtained at 1.0 *μ*M were used, in order to match the PKIS2 concentrations. PKIS1 contains 366 unique compounds with unique SMILES and ChEMBL IDs that were tested on 200 unique parent kinases having unique target ChEMBL IDs. When we include mutants/variants of the parent isoforms, there is a total of 224 targets with unique ChEMBL ASSAY IDs. Our processed PKIS1 data was therefore arranged as a matrix of 224 kinase targets by 366 compounds.

*PKIS2* The original PKIS2 was downloaded from https://doi.org/10.1371/journal.pone.0181585.s004. This set comprises 641 unique compound SMILES and 406 target columns. However, only of these 415 compounds were available to us from the original set for testing. We included only these compounds from the PKIS2 data set, so our bioactivity matrix has 406 targets by 415 compounds. PKIS2 activity values represent percent inhibition values observed at inhibitor concentrations of 1 *μ*M.

### Bioactivity-based prediction methodology

#### Setup

We let 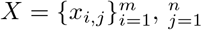 denote the bioactivity inhibition matrix that is available initially, where *m* denotes the number of kinase targets and *n* denotes the number of compounds. We use ***x**_i_*. for the vector of bioactivity results on target *i*, and ***x***_.*j*_ for column entries of this matrix (that is, bioactivity results for compound *j*). Let *I* = {1, 2, ⋯, *m*} denote the targets associated with rows of the data matrix *X*, and *J* = {1, 2, ⋯, *n*} denotes the set of available compounds.

For some methods, the kinase inhibition matrix *X* is reduced to a binary matrix 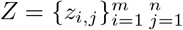, which captures empirical assessments of whether target *i* is inhibited (or not) by compound *j*. We use a target-wise threshold criterion (Eq. 1), based on the sample mean and sample standard deviation of each row ***x**_i_*., as follows:

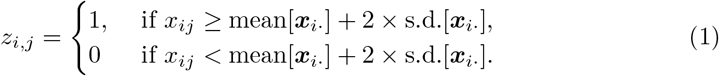

Our ultimate task is prediction from *X* of binary activities: *z_i*,j_*, on a new target *i*\* ∉ *I* for available compounds, *j* ∈ *J*. Our approach is first to identify from *X* a small *informer set* of compounds, *A* ⊂ *J*, on which bioactivity data *x_i_*,_j_* will be measured against target *i*\*. The data obtained from this experiment with the informer set, denoted by ***x***\* = {*x_i_*,_j_*, *j* ∈ *A*}, will be used to identify other compounds in the full compound set *J* that inhibit new target *i*\*, in the sense that *z_i_*,_j_* = 1. Machine-learning and statistical tools are used to design and study this approach, but except through general parallels with adaptive experimental design, the selection of an informer set is neither a supervised nor an unsupervised machine-learning task. We describe three novel heuristic methods that have favorable empirical characteristics: regression selection (RS), coding selection (CS), and adaptive selection (AS).

The informer-based ranking (IBR) methods that we propose entail partitions of the target set *I*, also referred to as a set of clusters, sometimes denoted by *S* = {*S*_1_, *S*_2_, ⋯, *S_K_*}. Methods differ as to how any candidate partition *S* is evaluated or acted upon. A partition *S* induces a labeling of targets *i* ∈ *I*, denoted *y* = (*y_i_*), where *y_i_* = *k* if (and only if) *i* ∈ *S_k_*. In all methods, the new target *i*\* becomes associated with one of the clusters by virtue of similarity of bioactivity profiles with other targets.

#### Regression Selection (RS)

The relationship between the training target space *I* and a new target (point) needs to be established in order to predict active compounds on novel kinase targets. The informer set serves to locate the new target in the training space. Unsupervised clustering is used to partition the target space; then the informer set is chosen from compounds that are predictive of cluster labels in a coupled, supervised analysis.

##### Informer set

The informer set is identified using clustering, regression, and feature selection. First, we classify the target space — the row space ***x**_i_*. — into clusters such that all targets within the same cluster exhibit a similar response to the compounds. For this task we considered `k-means`, which tries to minimize the sum of the within-cluster distances from each cluster centroid. Formally, given a parameter *K* as the number of clusters, and *m* data vectors ***x**_i_*.,…, ***x**_m_*., it aims to solve

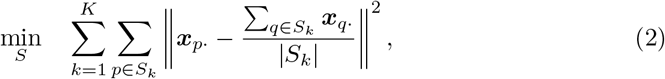

where *S* = {*S*_1_,…, *S_K_*} forms a partition of *I* = {1, ⋯, *m*}, and |*V*| denotes the cardinality of the set *V*. For robustness, we scale each column of the bioactivity data *X* linearly so that its entries lie in the range [0,1] prior to clustering analysis.

There are two difficulties in using *k*-means. The first is that its iterative process relies on random initialization, hence the results generally differ on each run. Secondly, we do not know in advance how to specify the number clusters *K*. To deal with the first problem, we use the `kmeans++` initialization procedure [32]. `kmeans++` guarantees that the expected final objective value is no more than *O*(log *K*) times larger than the optimal. To further improve robustness, we repeat the `kmeans++` procedure 100 times and choose the outcome that has the lowest objective value in Eq. 2. For the second issue of selecting *K*, we find the value that achieves the best performance in a five-fold cross-validation procedure.

Clusters serve to label the targets, as noted above. Namely, we set *y_i_* = *k* if *i* ∈ *S_k_* in Eq. 2. Next, multinomial logistic regression with a penalty term is applied to train a label classifier. In this approach, training data has the form {(***x**_i_*.,*y_i_*)}, over an appropriate set of targets *i*. The multi-class classifier is trained by fitting the multinomial logistic model. That is, we seek a set of coefficients, ***ω*** = {*ω*_10_, ⋯, *ω*_*K*0_, ***ω***_1_, ⋯, ***ω**_K_*}, by minimizing the objective function

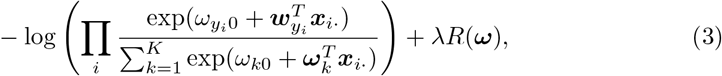

where the first term is the (negative) log-likelihood from the multinomial logistic model, *λ* ≥ 0 is a tuning parameter, and *R*(***ω***) is a penalty function. The coefficients ***ω**_k_*, one for each cluster, are vectors whose length equals the number of compounds, each of whose elements (***ω**_k_*)_*j*_ represents the weight that is applied to the activity measurement for compound j in predicting membership of cluster *k*. An appropriately chosen penalty yields sparse solutions in which coefficients of compounds that are only weakly predictive of the target cluster are set to zero. Since the whole coefficient vector ***ω**_.j_*:= ((***ω***_1_)_*j*_, (***ω***_2_)_*j*_, ⋯, (***ω_K_***)_*j*_) captures the influence of compound *j* on all targets, variable selection is achieved only if these *K* coefficients are shrunk simultaneously to zero. Therefore, we use the group LASSO norm [33] corresponding to the penalty function:

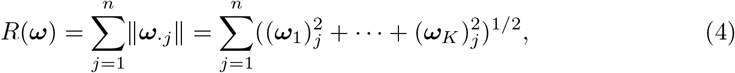

where 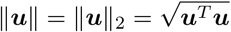 denotes the *L*_2_-norm.

We combine this regularized model with the greedy heuristic proposed in [34], which was shown to outperform the model obtained by directly solving Eq. 3 by choosing relevant features greedily, one at a time. This algorithm starts by setting *A* = Ϙ, then solves Eq. 3, and then selects the feature vector ***ω**_.j_* with the largest Euclidean norm, from among those features still represented in the regularization term *R*. It adds *j* to the informer set *A*, then re-solves Eq. 3 with all feature vectors ***ω**_.j_* for *j* ∈ *A* excluded from *R*. This process is repeated until either we have selected enough features (denoted by *n_A_*) or else the remaining ***ω**_.j_* included in the norm calculation are all zero. After selecting the features in this fashion, we retrain a model by solving Eq. 5 again using *only the selected features* and omitting the regularization term *R* altogether, that is,

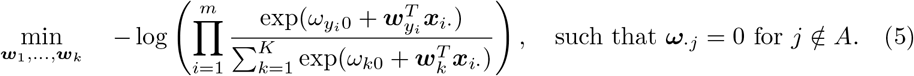

##### Compound ranking

Given any new target and its bioactivity with compounds from the informer set, we use the trained logistic regression model to predict the cluster label. With *y_i_*\* denoting the to-be-predicted cluster label of the new target *i*\* and ***x***\* the informer set compound activities for this target (i.e., the intermediate data), the multi-class logistic model asserts:

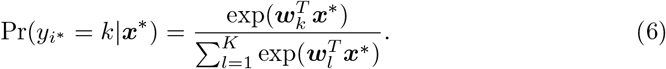

Using this probability and the centroids, we predict the whole activity vector of the new target with the vector of expected values:

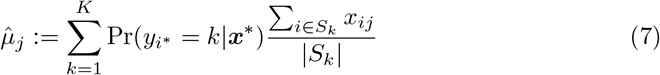

for compound *j*.

##### Parameter estimation

Given the whole active prediction procedure, we are able to conduct parameter selection using cross-validation. We conduct five-fold cross-validation to pick the best value of *K* for each evaluation metric and then retrain on the whole data matrix using the selected *K* to generate the final model for predicting the test data. Pseudo-code for the entire regression selection process is provided in Algorithm 1 in the Supplementary Information. We fix *λ* = 10^−6^ in Eq. 3 in our implementation.

#### Coding Selection (CS)

Coding selection works directly on the binary bioactivity data *Z* = {*z_i,j_*}_*i*∈*I,j*∈*J*_ rather than the quantitative inhibition measurements. The idea is to construct a single objective function to score potential informer compounds for how well they predict target activity the non-informer set. For a potential informer set *A* ⊂ *J*, CS considers that distinct rows of the sub-matrix *Z_A_* = {*z_i,j_*}_*i*∈*I*,*j*∈*A*_ constitute a kind of encoding of the kinase target space. Specifically, row *i* of *Z_A_*, which corresponds to kinase target *i*, is a length *n_A_* = 16 vector of zeros and ones. Among the 2^16^ = 65536 possible such vectors, only a relatively few distinct ones, numbering *L_A_* ≤ *n*, manifest themselves as rows of the sub-matrix in a given example. We call these distinct vectors *code words*, and denote them *q*_1_, *q*_2_, ⋯, *q_L_A__*. For some *K* ≤ *L_A_*, we introduce a partition *π* = {*b_k_*} of these code words, where each block holds a set of code words, and where *π* has *K* disjoint blocks. Together, the informer set A and the partition *π* induce a partition *S* = {*S*_1_, *S*_2_, ⋯, *S_K_*} of the targets *I* by the rule that *i* ∈ *S_k_* if (and only if) row *i* of the sub-matrix equals some code word *q_l_* ∈ *b_k_*. We emphasize that given candidates *A* and *π*, the target-space partition *S* is obtained using only information in the binary data sub-matrix *Z_A_*.

To provide some intuition for the coding construction, let’s look ahead to when intermediate data ***x***\* are obtained in experiments with informer set compounds *j* ∈ *A* on new target *i*\*. These inhibition measurements may also be binarized to produce bioactivity calls *z*\* = {*z_i*,j_*, *j* ∈ *A*}. If *z*\* exactly matches one of the code words *q_l_*, then any targets in *I* having this same code word are natural comparators for *i*\*. Their bioactivity profiles on the non-informer compounds may be the basis for a useful secondary prediction. In fact we may not have an exact match of the new code word, and there may be distinct code word profiles on the informer compounds that yield similar non-informer profiles. Therefore, we propose the following objective function to measure properties of the potential informer set *A* and the code-word partition *π* that are conducive to high-accuracy prediction on non-informer compounds:

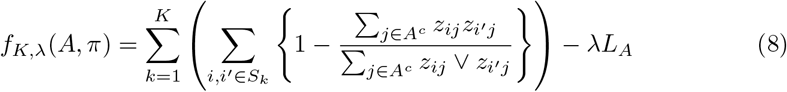

The inner summation in Eq. 8, which is over pairs of targets within cluster *S_k_*, accumulates pairwise differences between targets, as measured on the non-informer compounds, and using the *asymmetric binary* distance: among non-informer compounds that are active against either target, what fraction are not active against both? The outer sum is over clusters (partition blocks) of targets. The objective function value is low, therefore, if clusters induced by *A* and *π* and informer data are internally homogeneous on the non-informer data. For tuning parameter *λ* > 0, the penalty term *λL_A_* encourages informer sets that have many code words, in order to reduce the rate of extrapolation from intermediate data. The proposed scoring function (Eq. 8) is essentially non-parametric, allowing potentially complex relationships to exist between bioactivities of informer and non-informer compounds. This modeling flexibility comes at a cost, however, in that it is a combinatorial optimization task to identify the best *A* and *π* settings for any fixed *K* and *λ*.

Initially we sought to solve argmin *f_K,λ_*(*A, π*) approximately by Monte Carlo search. Fixing parameters *K* and *λ*, we randomly sample (*A*_1_, *π*_1_), (*A*_2_, *π*_2_), ⋯, (*A_B_, π_B_*) for a large number of trials *B*, such as 10^6^ or 10^7^. Each *A_b_* is a random subset of size *n_a_* = 16 taken from the full set of *n* compounds; then *π_b_* is a random partition of the code words from *Z_A_b__*. In numerical experiments, we found that marginally stabilizing compound scores is more effective than taking the informer set *Â* to be the sampled set *A_b_* having the lowest objective value (Eq 8). Specifically, we score every compound *j* ∈ *J* by

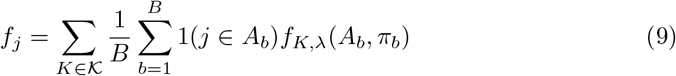

where 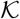 is a set of entertained cluster numbers. We used 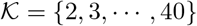, and fixed *λ* = 5 based on preliminary experimentation. The computed informer set *Â* contains the *n_A_* best (lowest) scoring compounds by this score.

##### Compound ranking

To proceed with ranking compounds, we require a code word *z*\* derived from the intermediate data *x*\* obtained on the new target *i*\*. A threshold level, such as used to binarize the original data (Eq. 1), may not be available. Instead we revert to the inhibition data on the informer compounds, say *X_Â_* (an *m* × *n_A_* sub-matrix of *X*), and we keep track of all the rows of *X_Â_* associated with each code word in the computed informer set. We compute a centroid for code-word *q_l_*, say, *c_l_*, by averaging the rows of *X_Â_* associated with *q_l_*. Then, the code-word centroid that is closest (in Euclidean distance) to the new data *x*\* is the derived code word *z*\* for target *i*\*.

Having our new target *i*\* provide code word *z*\* on the basis of intermediate data *x*\*, we next require a prediction of non-informer compounds that may also inhibit *i*\*. We score *j* ∈ *Â^c^* by their activity rates among the *n*\* targets with the same code word as *i*\*:

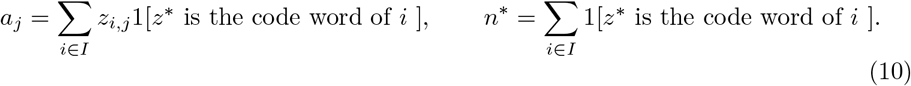

Our prediction of the active non-informer compounds is 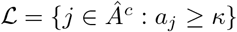 for some threshold *κ*. We set *κ* with an appeal to false-discovery-rate (FDR) control, recognizing that the Bernoulli trial *z_i*,j_* may be regarded as having success probability estimated by *a_j_*/*n*\*. Then a crude estimate of FDR of 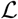 is

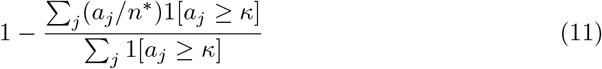

Similarly, we could estimate under the Bernoulli model the expected number of active non-informer compounds, Σ_*j*∈*A^c^*_ (*a_j_*/*n*\*), which may guide our choice of *κ*. Pseudo-code for the entire coding selection method is provided in Algorithm 2, Supplementary Information.

#### Adaptive Selection (AS)

The AS approach first identifies a base informer set of size *n*_0_ < *n_A_* compounds by a minor variation of the regression selection (RS) approach. We use *n*_0_ =8 and *n_A_* = 16. This step establishes both a clustering of the target space and the identity of *n*_0_ compounds that are predictive of the cluster labels. Next, AS adaptively grows the informer set, one compound at a time, so as to identify compounds that are predictive of non-informer bioactivity.

To identify the base informer set *A_o_*, the target space *I* is clustered using `k-means`, which aims to solve Eq. 2. With 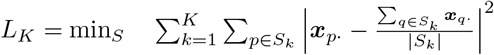, the number of clusters *K* is determined by

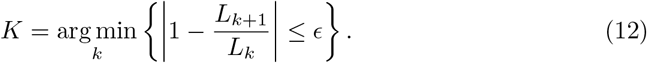

*ϵ* is a small value. In our calculations, *ϵ* = 0.02. Similar to RS, the clustered target space {*S_k_*} then serves as the response variable in a penalized multinomial regression to select the first *n*_0_ compounds of the informer set. Our specific implementation uses group LASSO as deployed in the `glmnet` R package for the multinomial response [35]. The regularization penalty is chosen so that precisely *n*_0_ compounds enter the predictive model.

For the remaining *n_A_* − *n*_0_ informer compounds, we augment the current set one compound at a time. Letting *A_c_* denote the current set, the next added compound solves:

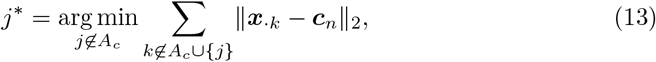

where ***x**_.k_* is the column vector in the inhibition matrix {*x_i,j_*}_*i*∈*i*,*j*∈*J*_ for compound *k*; 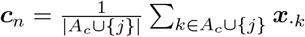 denotes the centroid of the current informer set. Eq. 13 finds the compound that minimizes the distance between informers and non-informers; the informer set is updated *A_c_* ← *A_c_* ∪ {*j*\*}. The final informer set is generated by iterating this process until there are *n_A_* compounds in *A_c_*.

For compound ranking, AS uses the same approach as CS (Eq. 11) after the code word *z*\* is acquired on generated informer set. Pseudo-code for AS is provided in Algorithm 3 of Supplementary Information.

#### Baseline Models (B)

As practical baseline approaches against which to compare our bioactivity-guided experimental design strategies (RS, CS, and AS), we applied two different informer selection methods: one based on compound structural diversity and the other leveraging the most frequent hitter (nonselective) kinase inhibitors as observed from the compound bioactivity matrix. Then, based on data returned from these informer selections, we applied three different chemical feature-based compound ranking methods, yielding a total of six strategies.

*Baseline informer set selection*. Baseline informer compounds were selected from each data matrix by one of two different methods:

- *Chemometric* selection (BC) – Compounds are grouped by scikit-learn’s hierarchical agglomerative clustering procedure (*n_A_* = 16 clusters, average linkage) using a Jaccard distance matrix computed from RDKit-derived Morgan chemical fingerprints (radius=2, 1024-bits) as features [36–38]. The 16 cluster medoids are taken as the informers.
- *Frequent Hitters* selection (BF) – Matrix compounds *j* are ranked in descending order by the number of targets on which each is labeled active, i.e., by *f_j_* =Σ*i*∈*I z_i,j_*, where *z_i,j_* indicates activity in the input bioactivity data (1). In other words, the informer set contains the 16 most broadly active compounds.

Note that in the cross validation study, the informer set needs to be recomputed each time target is left out.

*Baseline compound ranking*. After data are returned on the informer set, the remaining non-informer compounds were then ranked by three chemometric “hit expansion” methods:

- *Simple Expansion (s)* – ranks each non-informer by its distance to the nearest active informer compound as measured by Jaccard distance between Morgan fingerprints.
- *Loop Expansion (l)* – loops through active informer compounds in order of descending activity and prioritizes the nearest unranked non-informer compound to the current active informer based on fingerprint distance. The loop continues until all non-informers have been ranked.
- *Weighted Expansion (w)* – ranks each non-informer compound by Euclidean inner product of a target’s informer activity vector (16 normalized activities) and a compound’s vector of Jaccard similarities to those 16 informers. This scalar represents the “activity-weighted” similarity of each compound to the informer set. Compounds are prioritized in order of ascending values. Therefore, for target i*, we have a single activity vector ***x***\* comprising the 16 informer activities (normalized to [0,1]):

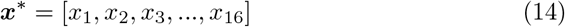 Each noninformer compound, *j*, has a similarity vector ***v*** representing the compound’s similarity to each of the informers tested on target *i*\*:

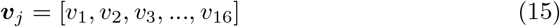 The weighted expansion score, *w*, for compound *j* on target *i*\* is then the Euclidean Inner product:

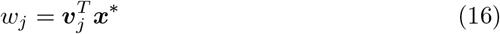

In the simple and loop expansion ranking methods, a binary label is required to designate “active” and “inactive” informers. This is problematic because this label depends on a target’s compound activity distribution (*μ*+2*σ* threshold), which is unknown prior to experimental screening of the compound set. To address this issue, we predict the activity threshold used for assigning a compound’s binary activity label on a given target using data returned on the 16 informer compounds. From these 16 informer activities, a threshold is inferred threshold from the available compound activity distributions on other targets in the matrix. This threshold for each target is the *kappa* parameter described above in the Coding Selection method.

#### Biological assays on novel microbial kinase targets

##### Mycobacterium tuberculosis PknB

Recombinant bacterial kinase (PknB) and bacterial substrate (GarA) were purified from *E. coli* following published procedures ([39]). The kinase inhibition assay was done using the Kinase Glo(R) kit from Promega similar to published procedures. PknB was added to plated kinase inhibitor libraries (the available compounds from PKIS 1 and 2) and incubated at room temperature for 10 minutes, after which ATP and GarA (protein substrate) were added. The final concentrations were: PknB 0.25 *μ*M, GarA 40 *μ*M, ATP 100 *μ*M, inhibitors 2 *μ*M, DMSO 1% in a final volume of 5 *μ*L. The kinase reaction proceeded at room temperature for 30 minutes and quenched by the addition of 5 *μ*L of Kinase Glo(R) reagent. The plate was allowed to develop for 10 minutes and luminescence was detected on a BMG PheraStar multiplate reader. Luminescence was converted to *μ*mol/minute of ATP consumed using a standard curve of ATP from 100 to 0 *μ*M. A negative control (no inhibitor) was used to determine percent activity. A positive control (GSK690693) was used to ensure a baseline and compare plate-to-plate variation. Data were analyzed using CDD Vault (Collaborative Drug Discovery, Inc.) to determine plate *Z*′ > 0.5 and report percent inhibition for each compound.

##### Epstein-Barr virus BGLF4

Viral kinase BGLF4 was graciously provided by the laboratory of Professsor Yongna Xing. BGLF4 was expressed with an N-terminal His_8_-MBP-dual-tag in insect cells, and purified over Ni^2+^-NTA resin (Qiagen) and then Maltose resin (Qiagen), followed by ion exchange chromatography (Source 15Q, GE Healthcare) and gel filtration chromatography (Superdex 200, GE Healthcare) to more than 95% homogeneity. The purified BGLF4 was then used for kinase inhibition assays using the C-terminal fragment peptide of retinoblastoma protein (RB) as substrate (Millipore Sigma cta# 12-439). The remaining assay parameters were the same as those applied for PknB except for the following changes. The final concentrations in the reaction medium were: BGLF4 0.004 *μ*g/*μ*L, RB 0.04 *μ*g/*μ*L, ATP 500 *μ*M, inhibitors 3 *μ*M, DMSO 0.3% in a final volume of 5 *μ*L. As a positive control, K252a (5 *μ*M) was used. The reaction proceeded at room temperature for 20 minutes and was then quenched by the addition of 5 *μ*L of Kinase Glo(R) reagent. ADP depletion proceeded for 40 minutes, followed by addition of 10 *μ*L of kinase detection reagent. The reactions were incubated for 1 hour prior to luminescence detection.

#### Model metrics and evaluation procedure

##### Metrics

To evaluate model performance, we applied three different virtual screening metrics, ROCAUC, NEF10, and FASR10. ROCAUC and NEF10 measure the extent to which a model prioritizes the active compounds in its ranking. ROCAUC is a standard metric in virtual screening [40] and applied generally in machine learning to evaluate classifiers. Enrichment Factor (EF) (Eq 17) is another commonly used metric for assessing virtual screening performance. EF reflects the fold increase in active compounds over that expected from random compound selection, for a subset of a compound library taken from some top ranking portion of a prioritized compound list.

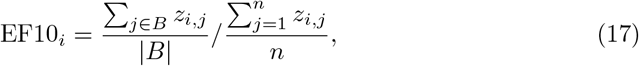

where *B* is the set of compounds among the top 10% of those ranked by a method applied to target *i*, and *z_i,j_* is as in (1).

However, the number of active compounds for each left-out target i varies from target to target (Fig S1). We apply a scaling scheme on EF at the top 10% (Eq 18), which enables better comparisons across targets exhibiting significant differences in active:inactive ratios.

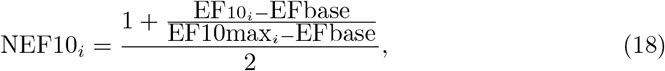

where EFbase is 1, which corresponds to random guessing; EF10max_*i*_ is the maximum theoretical EF10_*i*_, which means all actives are ranked at the top and depends on the number of actives for each target. Our NEF metric returns a value between 0.5 and 1, where a NEF10_*i*_ larger than 0.5 shows better ranking performance than random guessing-similar to ROCAUC. We selected the 10% threshold with consideration of the sizes of our informer (*n_A_* = 16) and full compound sets (*n* = 366 and *n* = 405). This threshold includes the 16 informers and 21 noninformer compounds in our PKIS1 evaluation.

For the ROCAUC and NEF10 metrics, experimental percent inhibition (activity) data were binarized using a target-specific *μ*+2*σ* threshold based on the activity distribution of the PKIS1 compounds for the kinase target. Actives were defined as compounds with greater than twice the standard deviations above the mean, as noted in (1). When applying the metrics, active informer compounds were counted as true positives, whereas inactive informers did not count against the models as false positives. It should be noted that the main purpose of the informer set is to facilitate accurate activity ranking on the non-informers. However, since informer compounds represent the highest priority compounds for testing, we reward models for retrieving active informers but refrain from penalizing models for choosing inactive informers. Some baseline models that rely upon binary compound labels occasionally failed to evaluate the noninformer compounds in cases where no active informers are returned. In such cases, metric scores reflecting random ranking were assigned to the model: ROCAUC and NEF10 of 0.5 and a FASR10 score of 0.0.

FASR10 assesses a model’s capacity to recognize different active chemotypes among the the top 10% of ranked compounds. The metric reflects the fraction of all active scaffolds identified on a given target within the compound set. Again, *z_i_* is the Boolean vector of compound binary activity labels on target *i* for compound set *J*. Let *O_J_* be the vector of chemical scaffold identifiers for compounds in J. The scaffold identifiers are arbitrary integer scaffold indices assigned to each of the generic Bemis-Murcko scaffold presented in *J*, as obtained using the `MurckoScaffold` module in RDKit [37,41]. Bemis-Murcko scaffolds were made generic by stripping hydrogens, converting all bonds to single, and setting all atom types to aliphatic carbon. The unique active scaffold identifiers are the set of all non-zero values in the Hadamard product vector:

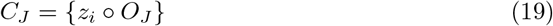

If we then let 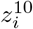 and 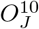 be the binary activity labels and scaffold IDs for the top 10% ranked compounds, the subset of unique active scaffolds recognized just among the top 10% of compounds is:

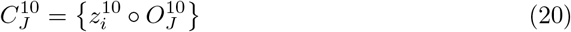

The fraction of active scaffolds recognized in the top 10% is:

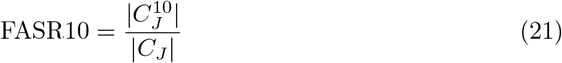

Note, active scaffolds were not considered *retrieved* unless an experimentally observed active member from that chemotype was in the top 10%. Cases arise where only inactive members of an active scaffold were obtained in the top 10% of the compound ranking. In such cases, the FASR10 metric does not count the chemotype as recognized.

Models are evaluated were first evaluated in a leave-one-target-out (LOTO) scheme: each of the 224 kinase targets i in PKIS1 data set is left out as a new target; one informer set *A_i_* and predictions on 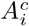 is generated using inhibition rate data of the other 223 targets.

##### Model evaluations

Performance of the models was evaluated in two stages. The first stage follows a retrospective leave-one-target-out (LOTO) evaluation scheme. Each kinase in the PKIS1 target set is removed and treated as a new target of interest. The PKIS1 compound activities are hidden for this target. An informer set is selected for this new target, the activities are revealed for the informers, and then the model rank orders the remaining noninformers using the informer data. The 9 models were evaluated in this stage using the 3 metrics described above. The second stage is a prospective evaluation of the 9 models as applied on two novel, non-human, kinase targets. In these evaluations, informer sets were generated twice for each model–once on each of the training matrices, PKIS1 and PKIS2. The remaining compounds (noninformers) from the corresponding matrix are then ranked on the two novel kinase targets using data returned for the informer sets and data within the corresponding PKIS1 or PKIS2 training matrix from which the informer set was selected. As in the retrospective PKIS1 LOTO evaluation, each model was assessed using the 3 metrics described above. However, in this prospective test on the new targets, each model was applied twice, using each of the PKIS data matrices, and therefore a total of 6 evaluations were performed on each model. We attempted to build a larger PKIS matrix by merging the PKIS1 and PKIS2 data matrices. The structure of the merged matrix, was however problematic in that the compound sets were nearly disjoint between PKIS1 and PKIS2. The resulting incomplete matrix lacks a structure that enables accurate imputation of the missing activity elements.

#### Code and data availability

PknB and BGLF4 screening data obtained at the UW-Carbone Cancer Center’s Small Molecule Screening Facility, formatted PKIS1 and PKIS2 datasets, and a Python implementation of the baseline IBR methods, evaluation metrics, and plotting procedures are available here: https://github.com/SpencerEricksen/informers. Matlab code and documentation involving the *RS* method is available here: https://github.com/leepei/informer. An R package for running *CS* and *AS* methods is available here: https://github.com/wiscstatman/esdd/tree/master/informRset.

## Acknowledgments

This work was funded by University of Wisconsin-Madison Carbone Cancer Center Support Grant P30 CA014520, NIH 1U54AI117924 to the UW Center for Predictive Computational Phenotyping, NSF awards 1321762 and 1148698 to the UW Center for High-Throughput Computing and Open Science Grid, and the UW-Madison Office of the Vice Chancellor for Research and Graduate Education with funding from the Wisconsin Alumni Research Foundation. We thank Rob Nowak and Sebastian Raschka for feedback on the manuscript. We also thank Yongna Xing and Vitali Stanevich for providing BGLF4 protein for assays and feedback on the manuscript.

## Supporting Information

### Figures and Tables

**S1 Table. Informer selections among PKIS1 compound set by IBR method**. From 366 PKIS1 matrix compounds, 16 informer compounds were selected each IBR method. The union of all informer compounds across 5 IBRs is listed. Informers selected by each method are indicated black dots. Informer activities are reported as normalized percent inhibition values [0,1]. Informers considered active have activities reported in boldface.

**S2 Table. Metrics evaluations against two new kinase targets (a) PKN and (b) BGLF4 using PKIS1 or PKIS2 matrices**. IBR strategies were also compared against novel kinase targets, (a) Pknb and (b) BGLF4, which do not belong to either of the PKIS1 and PKIS2 target sets.

**S3 Table. PKIS1 Leave-One-Target-Out cross validation performance c metrics (a) ROCAUC, (b) NEF10, and (c) FASR10**. Some baseline models rely on binary labels for informers (BC_s_, BC_l_, BF_s_, and BF_l_), which are essentially expansion strategies, exhibit a fairly significant failure rate in our PKIS LOTO validation. These methods cannot proceed with non-informer ranking in cases where active informers are returned. For performance evaluations in the main text, we considered these cases as methods’ failures on a target and assigned the method-target combination a performance metric reflecting a random ranking (e.g., ROC 0.5). Failure rates were particularly high among those using chemometric informer selection method (BC_s_, BC_l_), with no actives returned, and hence failure, on 68 of the 224 PKIS1 targets (30.4%). As expected, our frequent hitters informer selection method (BF_s_, BF_l_) has a relatively lower failure rate with failure on 12 targets (5.4%). Given the high hit rates for compounds within the PKIS1 matrix (5.6 +/− 2.1% using target-specific *μ*+2*σ* threshold), we anticipate that failure rates could be get much higher as we extend our strategies to chemogenomic matrices with broader target sets, which are likely to be more sparse. These issues are circumvented by ranking strategies that accommodate continuous activity data and thus proceed with inferences based on weakly active informers, rather than failing when no informers meet the active binary threshold. Thus, the baseline *w* ranking strategies and all of the purely bioactivity based IBR strategies were not subject to this type of failure.

**Table S1.** Informer selections among PKIS1 compound set by IBR method. From 366 PKIS1 matrix compounds, 16 informer compounds were selected by each IBR method. The union of informers across 5 IBRs is listed. Informers selected by each method are indicated black dots. Informer activities are reported as normalized percent inhibition values [0,1]. Activities of informers considered active are reported in boldface.

| PubChem CID | Activity |  | IBR methods |  |  |  |  |
| --- | --- | --- | --- | --- | --- | --- | --- |
|  | PknB | BGLF4 | BC<sub>w</sub> | BF<sub>w</sub> | RS | CS | AS |
| 10173796 | **0.42** | 0.07 |  | • |  | • | • |
| 10308522 | 0.00 | 0.05 |  | • |  | • |  |
| 11163861 | 0.11 | 0.15 |  | • |  | • |  |
| 11281930 | 0.00 | **0.39** |  |  | • |  |  |
| 11328632 | 0.00 | 0.00 |  |  | • |  |  |
| 11626927 | 0.00 | 0.08 |  |  | • | • |  |
| 11751266 | 0.00 | 0.02 |  |  | • |  |  |
| 16048303 | 0.03 | 0.00 | • |  |  |  |  |
| 23646938 | 0.01 | 0.07 |  |  |  | • | • |
| 25023715 | 0.00 | 0.02 |  |  | • |  | • |
| 25211574 | 0.00 | **0.29** |  |  | • |  | • |
| 25218584 | 0.02 | 0.00 |  |  | • |  |  |
| 25218593 | 0.00 | 0.00 |  |  | • |  |  |
| 25218600 | 0.00 | **0.63** |  | • |  | • | • |
| 25218601 | 0.00 | 0.00 |  | • |  | • | • |
| 25218614 | 0.00 | **0.70** | • | • |  | • |  |
| 44397010 | 0.00 | 0.00 | • |  | • |  |  |
| 44397250 | 0.04 | 0.00 |  |  | • |  |  |
| 44397453 | 0.06 | 0.05 |  |  | • |  |  |
| 44418539 | 0.01 | 0.00 | • |  |  |  |  |
| 44532204 | 0.00 | 0.01 | • |  |  |  | • |
| 44532523 | 0.01 | 0.00 | • |  |  |  |  |
| 44536036 | 0.01 | 0.00 |  | • |  | • |  |
| 44581245 | 0.00 | 0.00 |  |  |  |  | • |
| 448008 | 0.00 | 0.00 | • |  |  |  |  |
| 53239967 | 0.00 | **0.46** |  | • |  | • | • |
| 5329829 | 0.00 | 0.00 |  | • | • |  | • |
| 5329854 | 0.00 | 0.09 | • |  |  |  |  |
| 5482344 | 0.00 | **0.73** |  | • |  | • |  |
| 56604013 | **0.42** | 0.00 |  |  |  |  | • |
| 56604034 | **0.27** | 0.00 |  | • |  | • | • |
| 57391096 | 0.01 | 0.00 |  | • | • | • | • |
| 57396353 | 0.04 | 0.11 |  | • |  |  |  |
| 57399798 | 0.01 | 0.02 | • |  |  |  |  |
| 6539047 | 0.00 | 0.00 | • |  |  |  |  |
| 6539056 | 0.00 | 0.00 |  |  | • |  |  |
| 6539081 | 0.07 | **0.33** |  |  | • | • | • |
| 6539107 | 0.00 | **0.46** |  | • |  | • |  |
| 6539108 | 0.03 | **0.46** |  | • |  | • |  |
| 6539361 | 0.00 | 0.03 | • |  |  |  |  |
| 6539593 | 0.00 | 0.00 | • |  |  |  |  |
| 766949 | 0.03 | 0.02 | • |  |  |  |  |
| 9604911 | 0.00 | 0.00 |  |  |  |  | • |
| 9822610 | 0.00 | 0.00 |  | • |  |  |  |
| 9826308 | 0.03 | 0.00 | • |  |  |  |  |
| 9901964 | 0.00 | 0.00 | • |  |  |  |  |
| 9910722 | 0.00 | 0.00 | • |  |  |  |  |
| 9927432 | 0.00 | 0.06 |  |  | • |  | • |

**Table S2.**
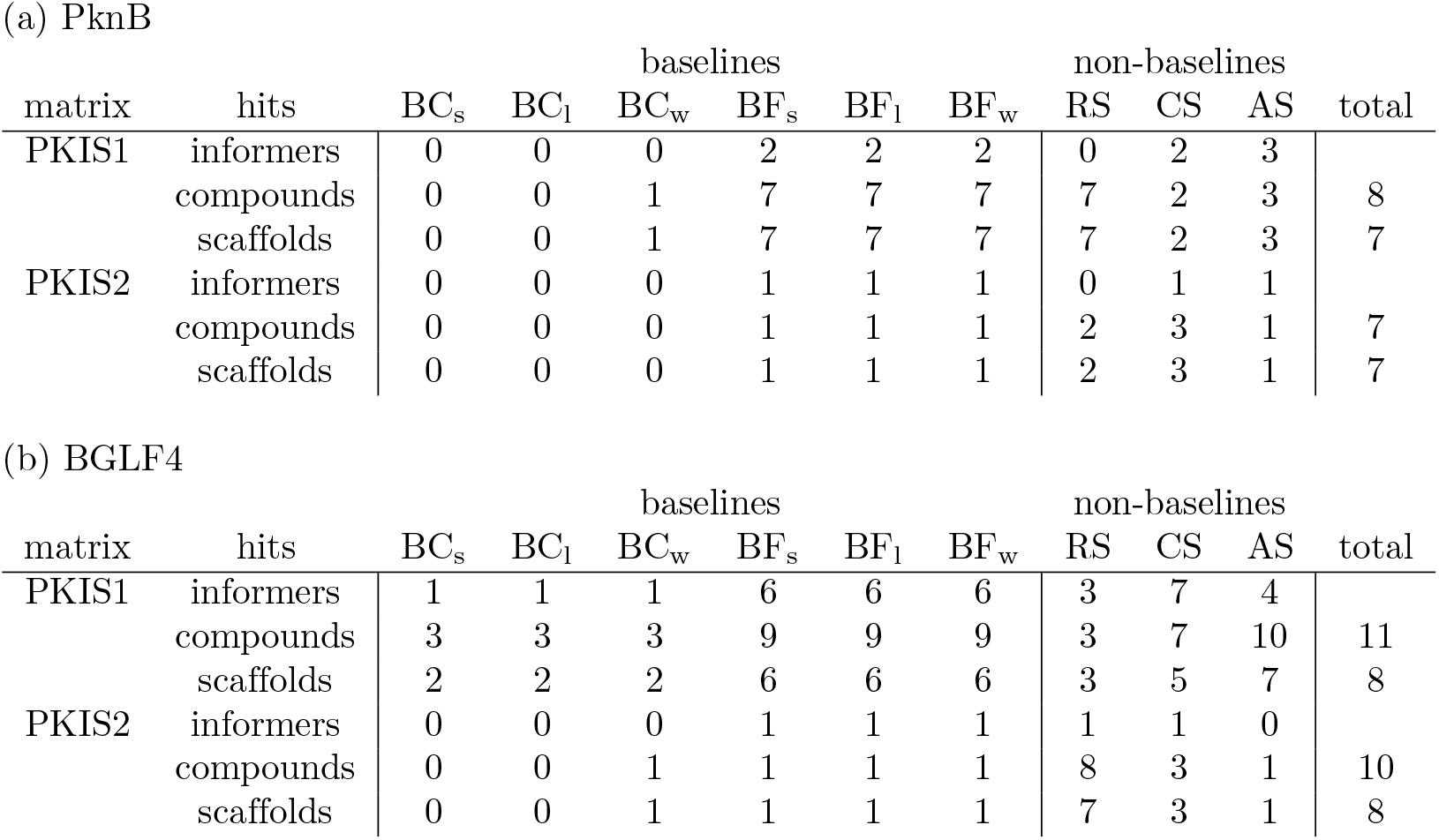
Retrieval counts by the various methods on new kinase targets (a) PknB and (b) BGLF4 using PKIS1 or PKIS2 matrices. The values below each of the IBR methods indicate the number of active informers observed (out of 16), the number of active compounds identified in the top 10% ranking compounds by each method, and the number of active scaffolds recognized in the top 10%. The total number of experimentally determined compounds and scaffolds is indicated in the *total* column.

| matrix | hits | baselines |  |  |  |  |  | non-baselines |  |  | total |
| --- | --- | --- | --- | --- | --- | --- | --- | --- | --- | --- | --- |
|  |  | BC<sub>s</sub> | BC<sub>l</sub> | BC<sub>w</sub> | BF<sub>s</sub> | BF<sub>l</sub> | BF<sub>w</sub> | RS | CS | AS |  |
| PKIS1 | informers | 0 | 0 | 0 | 2 | 2 | 2 | 0 | 2 | 3 |  |
|  | compounds | 0 | 0 | 1 | 7 | 7 | 7 | 7 | 2 | 3 | 8 |
|  | scaffolds | 0 | 0 | 1 | 7 | 7 | 7 | 7 | 2 | 3 | 7 |
| PKIS2 | informers | 0 | 0 | 0 | 1 | 1 | 1 | 0 | 1 | 1 |  |
|  | compounds | 0 | 0 | 0 | 1 | 1 | 1 | 2 | 3 | 1 | 7 |
|  | scaffolds | 0 | 0 | 0 | 1 | 1 | 1 | 2 | 3 | 1 | 7 |

(b) BGLF4
| matrix | hits | baselines |  |  |  |  |  | non-baselines |  |  | total |
| --- | --- | --- | --- | --- | --- | --- | --- | --- | --- | --- | --- |
|  |  | BC<sub>s</sub> | BC<sub>l</sub> | BC<sub>w</sub> | BF<sub>s</sub> | BF<sub>l</sub> | BF<sub>w</sub> | RS | CS | AS |  |
| PKIS1 | informers | 1 | 1 | 1 | 6 | 6 | 6 | 3 | 7 | 4 |  |
|  | compounds | 3 | 3 | 3 | 9 | 9 | 9 | 3 | 7 | 10 | 11 |
|  | scaffolds | 2 | 2 | 2 | 6 | 6 | 6 | 3 | 5 | 7 | 8 |
| PKIS2 | informers | 0 | 0 | 0 | 1 | 1 | 1 | 1 | 1 | 0 |  |
|  | compounds | 0 | 0 | 1 | 1 | 1 | 1 | 8 | 3 | 1 | 10 |
|  | scaffolds | 0 | 0 | 1 | 1 | 1 | 1 | 7 | 3 | 1 | 8 |

**Table S3.** Metrics evaluations against two new kinase targets (a) PKNB and (b) BGLF4 using PKIS1 or PKIS2 matrices. IBR strategies applied prospectively on novel kinase targets, (a) Pknb and (b) BGLF4, which do not belong to either of the PKIS1 and PKIS2 target sets.

| matrix | metric | baselines |  |  |  |  |  | non-baselines |  |  |
| --- | --- | --- | --- | --- | --- | --- | --- | --- | --- | --- |
|  |  | BC<sub>s</sub> | BC<sub>l</sub> | BC<sub>w</sub> | BF<sub>s</sub> | BF<sub>l</sub> | BF<sub>w</sub> | RS | CS | AS |
| PKIS1 | ROCAUC | – | – | 0.33 | 0.95 | 0.96 | 0.91 | 0.89 | 0.70 | 0.68 |
|  | NEF10 | – | – | 0.51 | 0.93 | 0.93 | 0.93 | 0.93 | 0.58 | 0.65 |
|  | FASR10 | 0.00 | 0.00 | 0.12 | 0.88 | 0.88 | 0.88 | 0.88 | 0.25 | 0.38 |
| PKIS2 | ROCAUC | – | – | 0.32 | 0.47 | 0.47 | 0.45 | 0.79 | 0.51 | 0.65 |
|  | NEF10 | – | – | 0.44 | 0.52 | 0.52 | 0.52 | 0.60 | 0.68 | 0.52 |
|  | FASR10 | 0.00 | 0.00 | 0.00 | 0.14 | 0.14 | 0.14 | 0.29 | 0.43 | 0.14 |

(b) BGLF4
| matrix | metric | baselines |  |  |  |  |  | non-baselines |  |  |
| --- | --- | --- | --- | --- | --- | --- | --- | --- | --- | --- |
|  |  | BC<sub>s</sub> | BC<sub>l</sub> | BC<sub>w</sub> | BF<sub>s</sub> | BF<sub>l</sub> | BF<sub>w</sub> | RS | CS | AS |
| PKIS1 | ROCAUC | 0.38 | 0.38 | 0.35 | 0.94 | 0.96 | 0.89 | 0.74 | 0.78 | 0.94 |
|  | NEF10 | 0.60 | 0.60 | 0.60 | 0.90 | 0.85 | 0.90 | 0.60 | 0.80 | 0.95 |
|  | FASR10 | 0.25 | 0.25 | 0.25 | 0.75 | 0.75 | 0.75 | 0.38 | 0.62 | 0.88 |
| PKIS2 | ROCAUC | – | – | 0.52 | – | – | 0.58 | 0.88 | 0.57 | 0.75 |
|  | NEF10 | – | – | 0.50 | – | – | 0.50 | 0.89 | 0.61 | 0.50 |
|  | FASR10 | 0.00 | 0.00 | 0.12 | 0.12 | 0.12 | 0.12 | 0.88 | 0.38 | 0.12 |

**Table S4.** (a) ROCAUC, (b) NEF10, and (c) FASR10 in Leave-One-Target-Out Cross Validation on PKIS1. Nine IBR methods were evaluated on 224 PKIS1 targets using standard VS metrics that reflect active retrieval, ROCAUC and NEF10. FASR10 was also evaluated to reflect the chemical diversity of the actives retrieved.

| Methods | baselines |  |  |  |  |  | non-baselines |  |  |
| --- | --- | --- | --- | --- | --- | --- | --- | --- | --- |
|  | BC<sub>s</sub> | BC<sub>l</sub> | BC<sub>w</sub> | BF<sub>s</sub> | BF<sub>l</sub> | BF<sub>w</sub> | RS | CS | AS |
| mean | 0.62 | 0.62 | 0.63 | 0.79 | 0.79 | 0.79 | 0.90 | 0.81 | 0.84 |
| median | 0.56 | 0.56 | 0.67 | 0.83 | 0.82 | 0.81 | 0.92 | 0.83 | 0.88 |
| stdev | 0.16 | 0.15 | 0.22 | 0.14 | 0.14 | 0.13 | 0.11 | 0.14 | 0.13 |

(b)NEF10
| Methods | baselines |  |  |  |  |  | non-baselines |  |  |
| --- | --- | --- | --- | --- | --- | --- | --- | --- | --- |
|  | BC<sub>s</sub> | BC<sub>l</sub> | BC<sub>w</sub> | BF<sub>s</sub> | BF<sub>l</sub> | BF<sub>w</sub> | RS | CS | AS |
| mean | 0.61 | 0.61 | 0.62 | 0.74 | 0.74 | 0.74 | 0.80 | 0.79 | 0.82 |
| median | 0.57 | 0.57 | 0.60 | 0.74 | 0.74 | 0.72 | 0.82 | 0.79 | 0.85 |
| stdev | 0.12 | 0.12 | 0.13 | 0.13 | 0.13 | 0.13 | 0.13 | 0.14 | 0.13 |

(c)FASR10
| Methods | baselines |  |  |  |  |  | non-baselines |  |  |
| --- | --- | --- | --- | --- | --- | --- | --- | --- | --- |
|  | BC<sub>s</sub> | BC<sub>l</sub> | BC<sub>w</sub> | BF<sub>s</sub> | BF<sub>l</sub> | BF<sub>w</sub> | RS | CS | AS |
| mean | 0.24 | 0.24 | 0.31 | 0.51 | 0.52 | 0.52 | 0.68 | 0.65 | 0.71 |
| median | 0.21 | 0.21 | 0.29 | 0.50 | 0.53 | 0.50 | 0.72 | 0.65 | 0.75 |
| stdev | 0.22 | 0.22 | 0.21 | 0.23 | 0.22 | 0.21 | 0.22 | 0.26 | 0.23 |

**Table S5.**
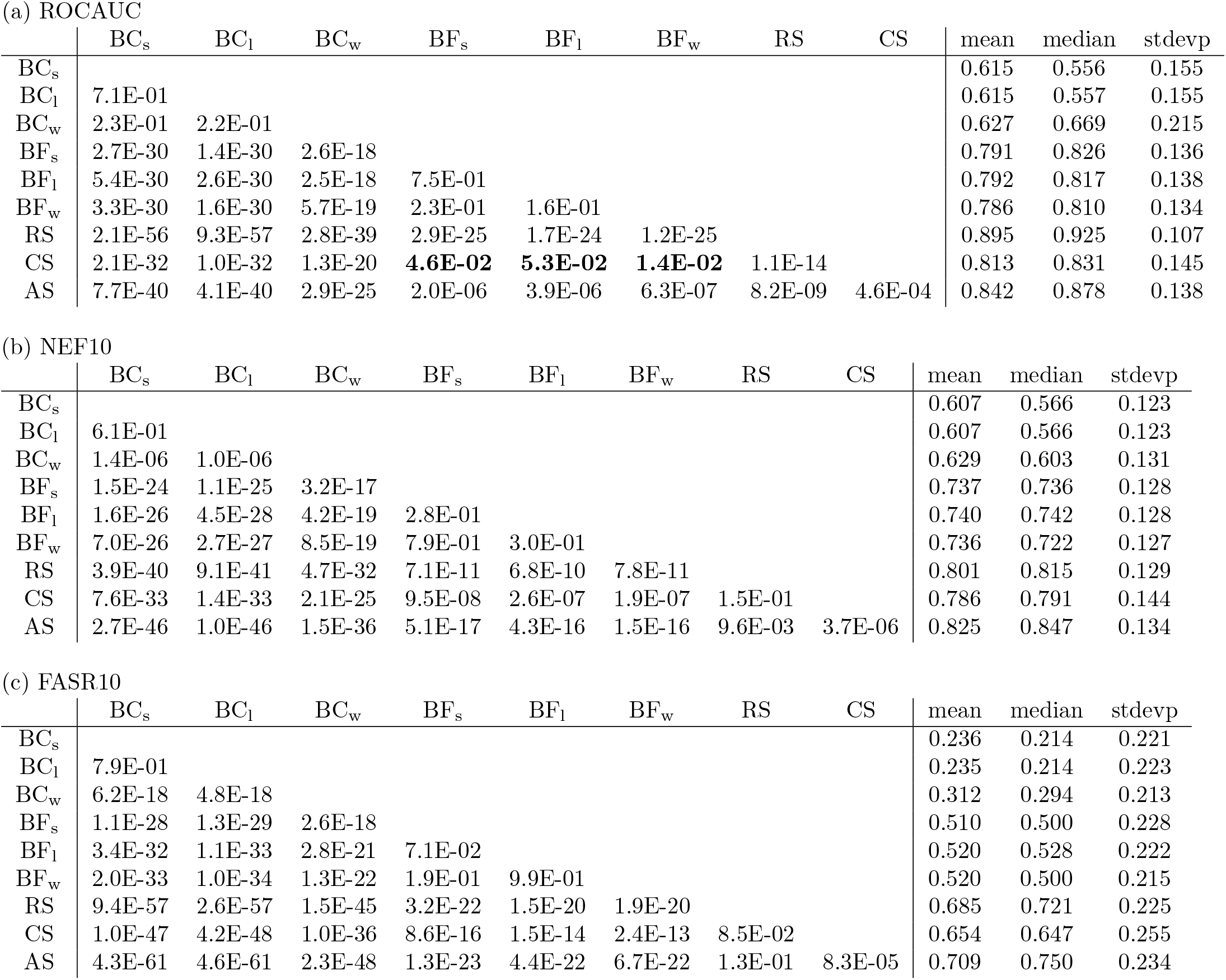
*P*-values in methods comparison for PKIS1 LOTO VS. VS metrics (a) ROCAUC, (b) NEF10, and (c) FASR10 in Leave-One-Target-Out Cross Validation on PKIS1 are compared across 9 IBR methods evaluated on 224 PKIS1 targets. Bold font is used to indicate *p*-values that fail to pass *α* = 0.05 threshold for significance between baseline methods. To collectively compare all 6 baseline IBRs against each non-baseline IBR, we imposed a Šidák multiple comparison correction with 6 hypotheses. This increases the stringency of the statistical threshold applied on each of the 6 individual tests to *α* = 8.5*E*3. However, after applying this correction, non-baseline methods remained statistically superior to all baselines for all metrics except for CS when considering the ROCAUC metric.

**Fig S1.**
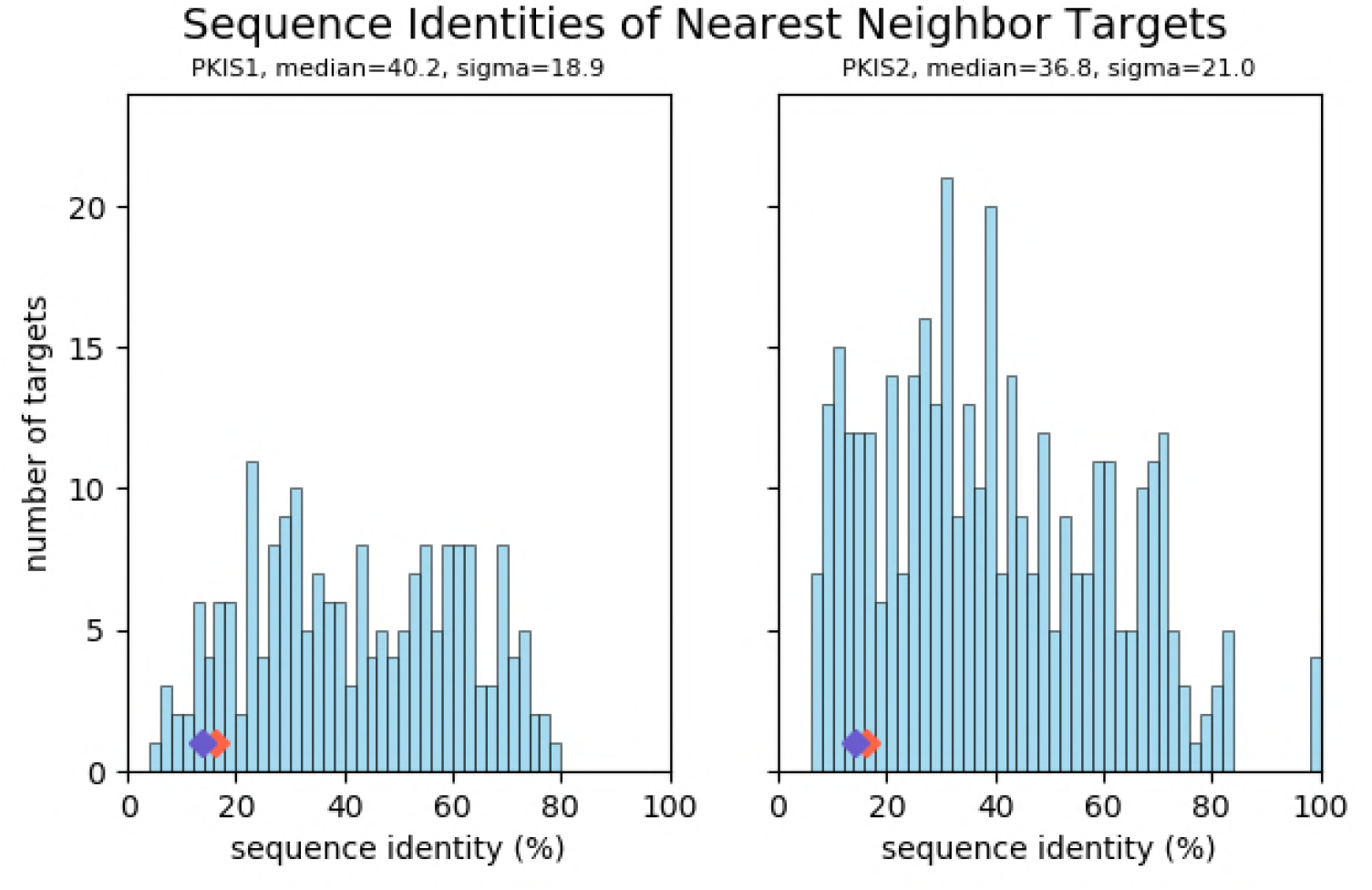
Nearest-neighbor sequence identity distributions for PKIS kinases. A sequence similarity matrix (% sequence identity of kinase domains) was determined for most members of the PKIS1 and PKIS2 kinase sets (mutants removed). The sequences of targets BGLF4 and PknB were also included. The histograms show the distribution of nearest-neighbor sequence identities among matrix kinases (PKIS1 or PKIS2). The blue (BGLF4) and red (PknB) diamonds indicate nearest neighbor sequence identities observed for these targets in the PKIS1 and PKIS2 kinase sets. BGLF4 and PknB do not have closely related neighbors in the sets.

**Fig S2.**
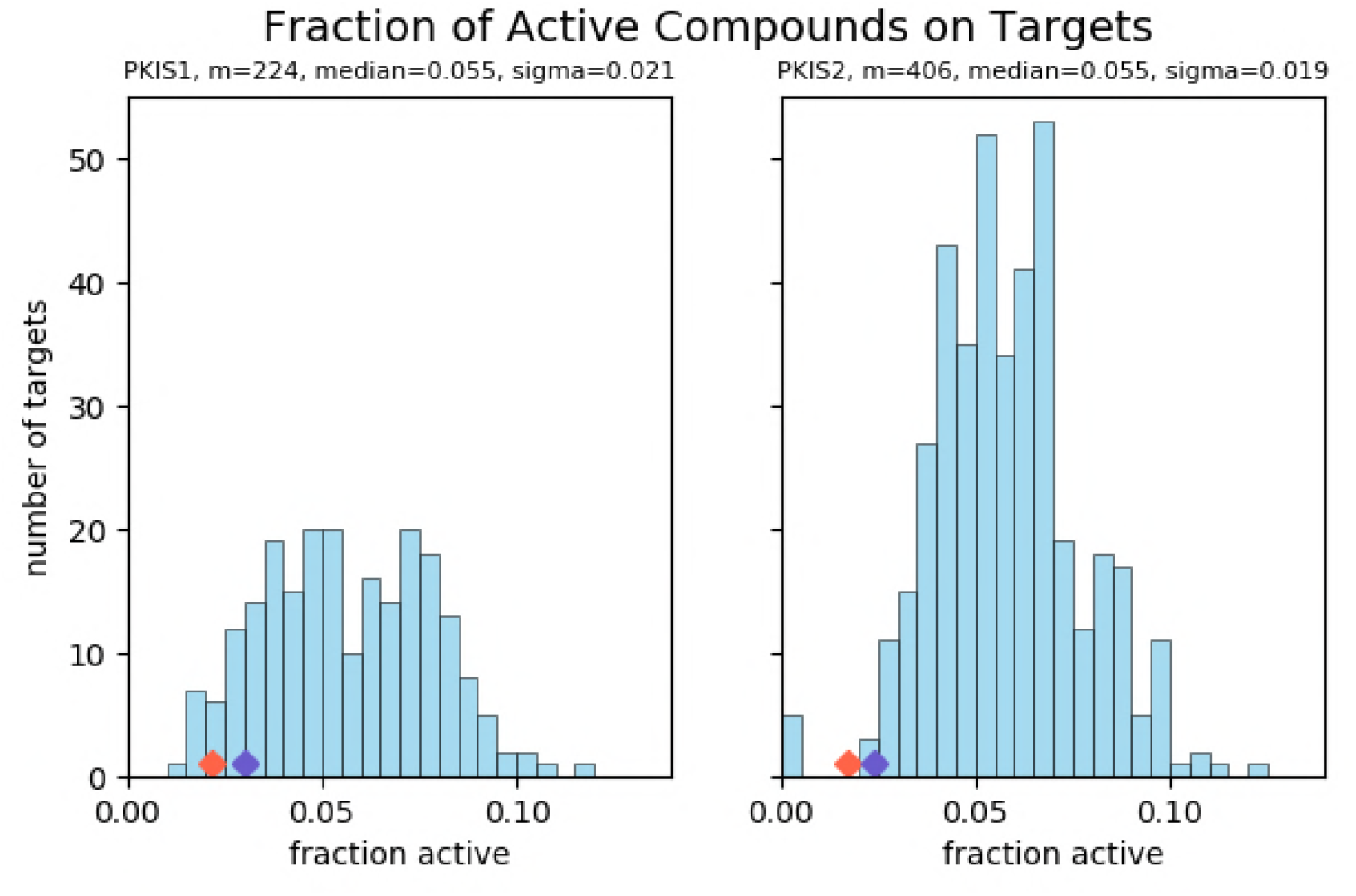
The distribution of active compound fractions across kinase targets in PKIS1 and PKIS2. The variation in the class imbalance is even wider when a universal threshold is applied (percent inhibition) over all targets. Diamonds indicate the fraction of active compounds for BGLF4 (blue) and PknB (red) in the PKIS1 and PKIS2 compound sets.

**Fig S3.**
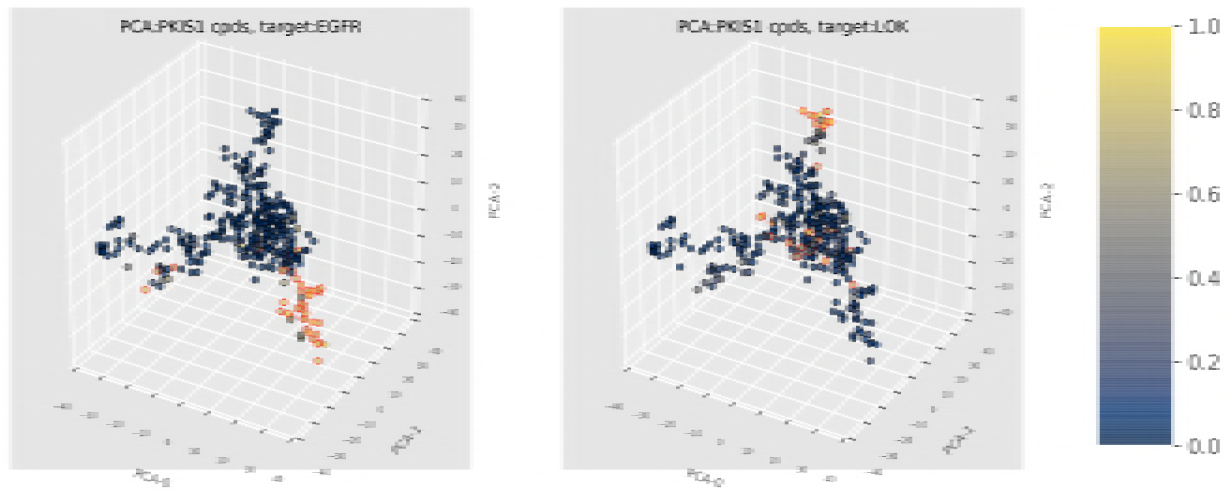
Projection of PKIS1 compounds along the 3 primary components taken from PCA on their Morgan fingerprints. Target kinases EGFR and LOK are shown here as examples. PKIS1 compounds (points) are colored according to their experimental activity on the target: yellow indicates high activity (strong inhibition) and blue is low. Active compounds (exceeding threshold) have markers outlined in red. On these example kinase targets, separated regions of active chemical space can be observed.

**Fig S4.**
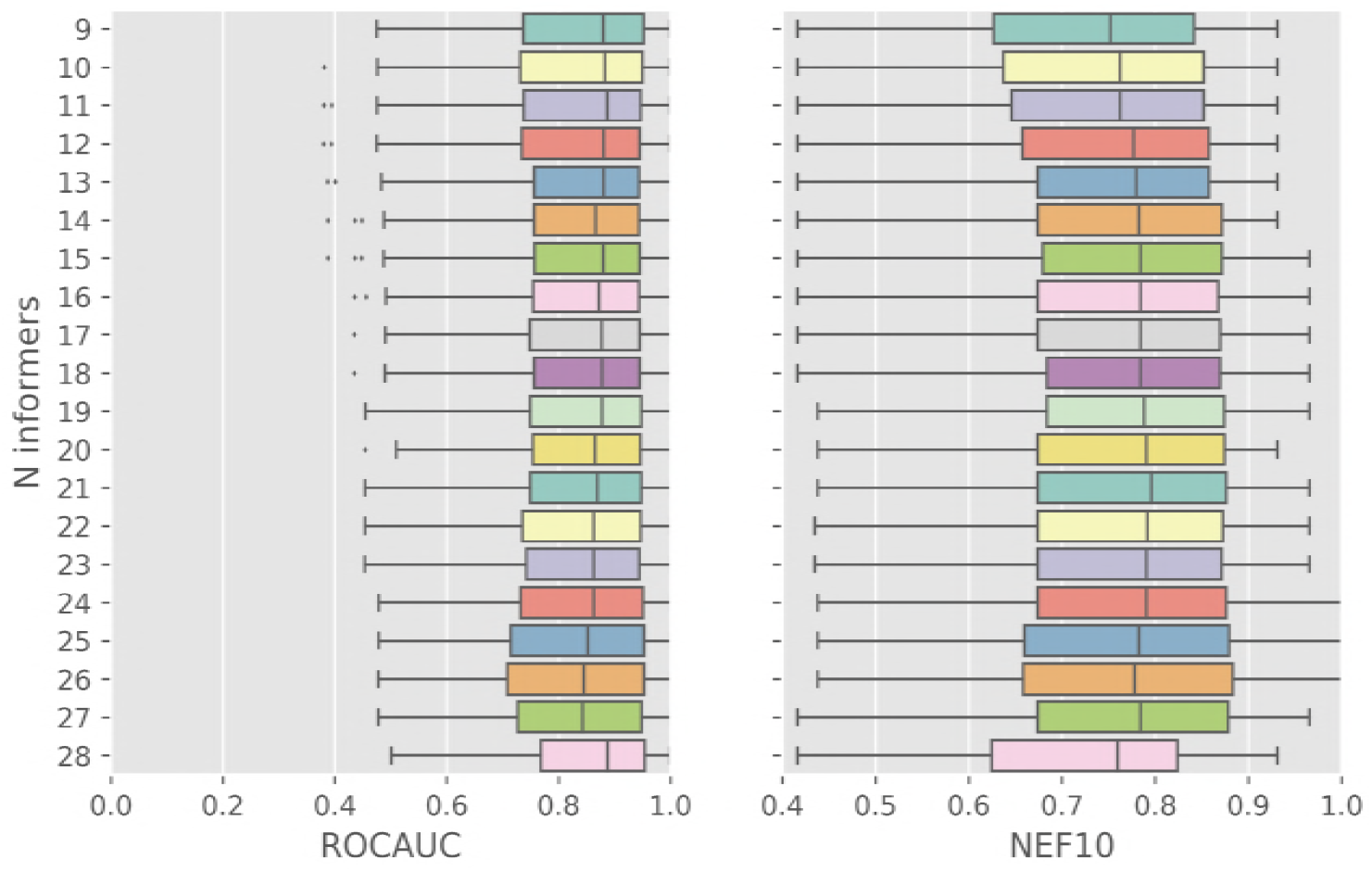
PKIS1 LOTO VS performance of AS method as function of informer set size. We examined the relationship between informer set size (*n_A_* =9 to 28) for IBR method AS and virtual screening performance in terms of ROCAUC and NEF10 metrics.

**Fig S5.**
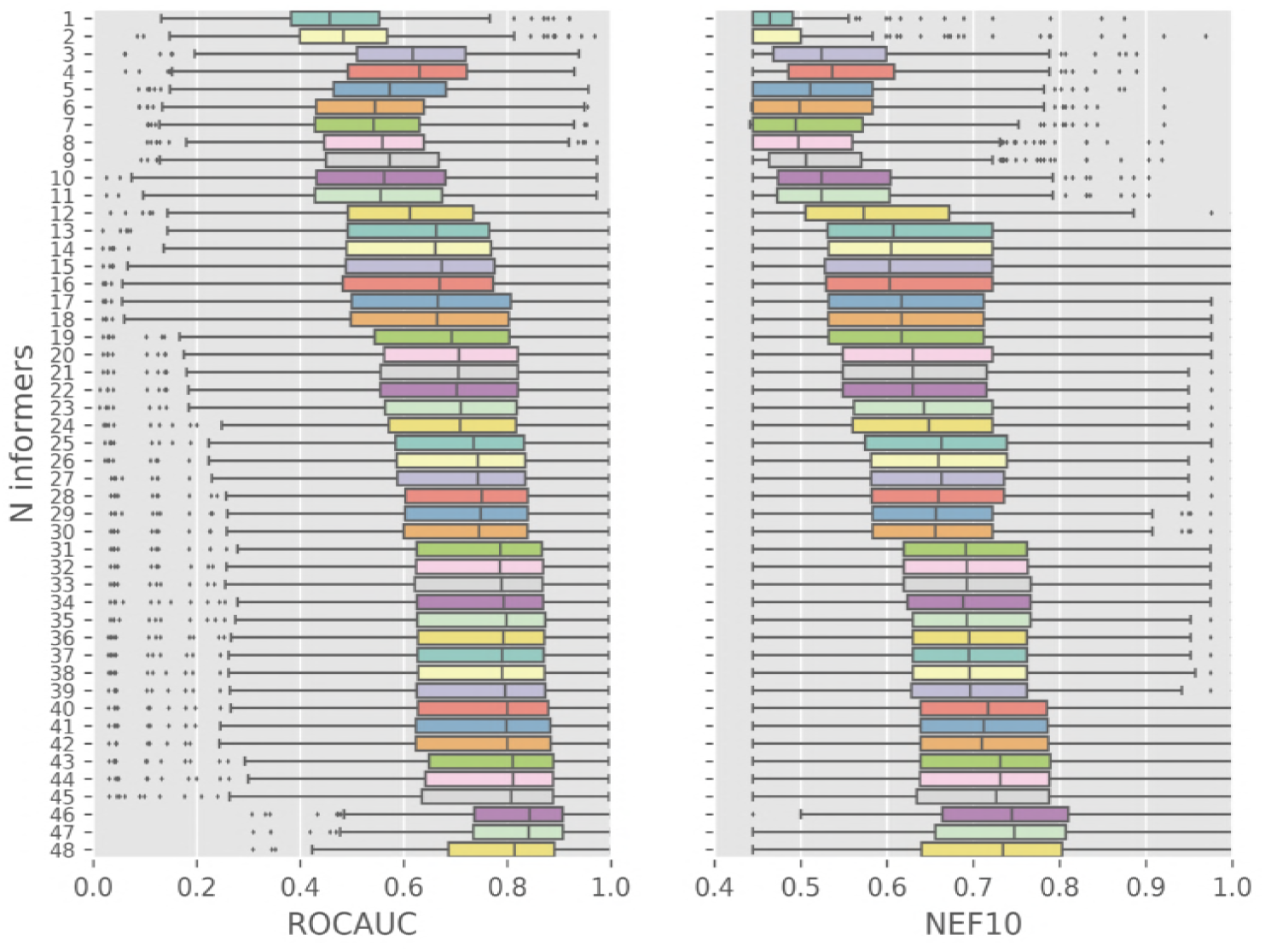
PKIS1 LOTO virtual screening performance as a function of informer set size for baseline method BC_w_, ROCAUC (left) and NEF10 (right).

**Fig S6.**
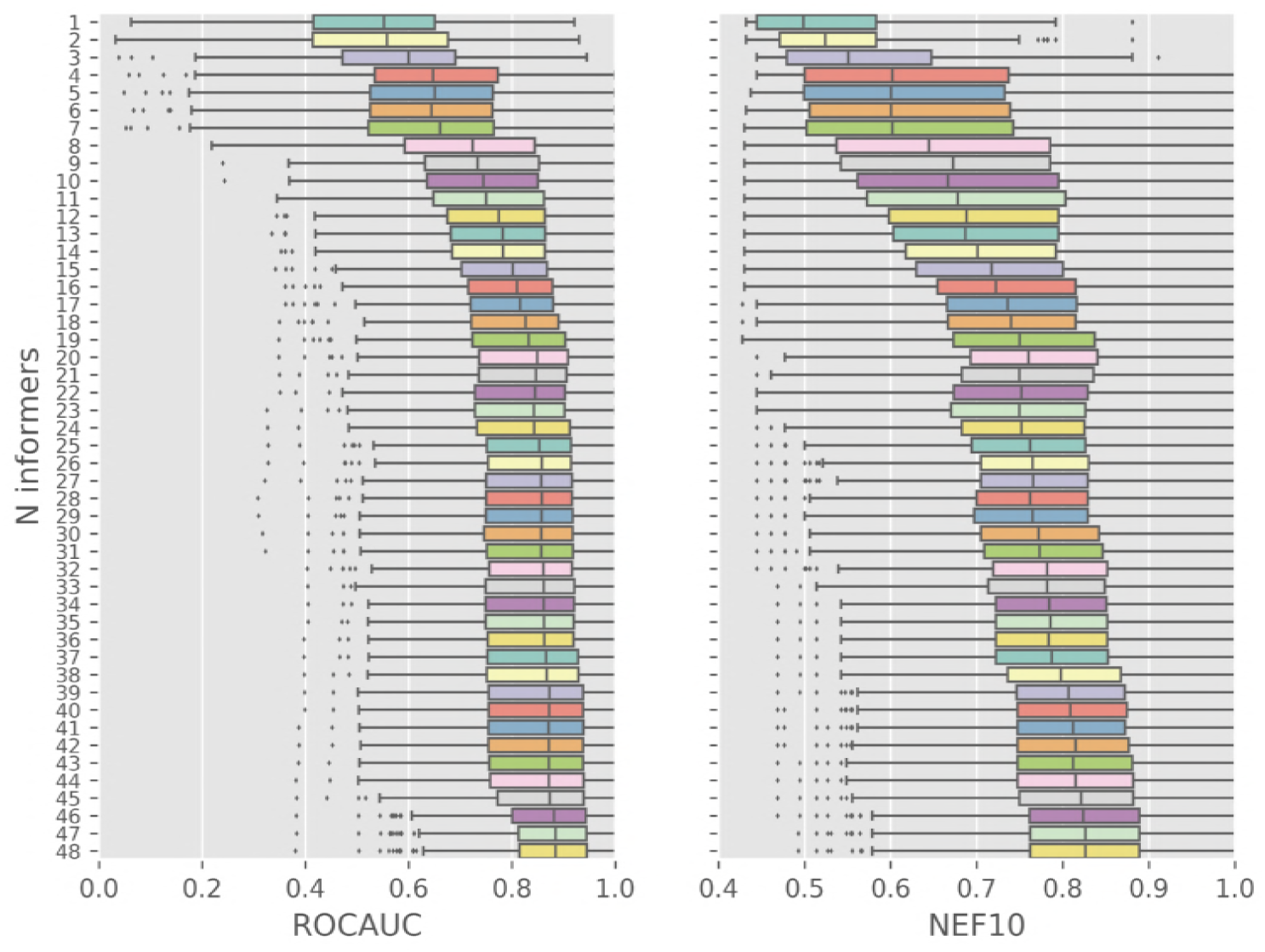
PKIS1 LOTO virtual screening performance as a function of informer set size for baseline method BF_w_, ROCAUC (left) and NEF10 (right).

## Algorithm pseudo-code 103

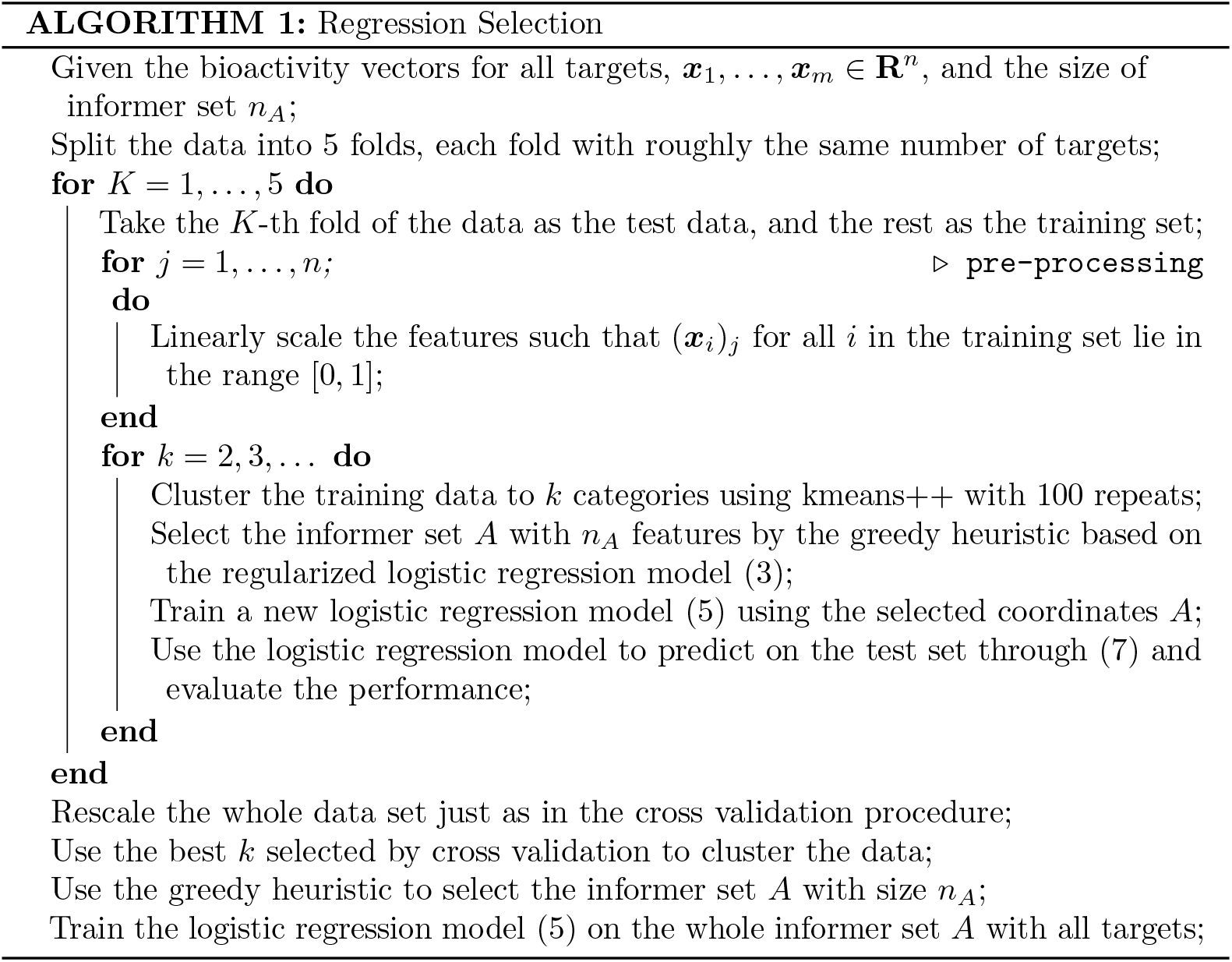

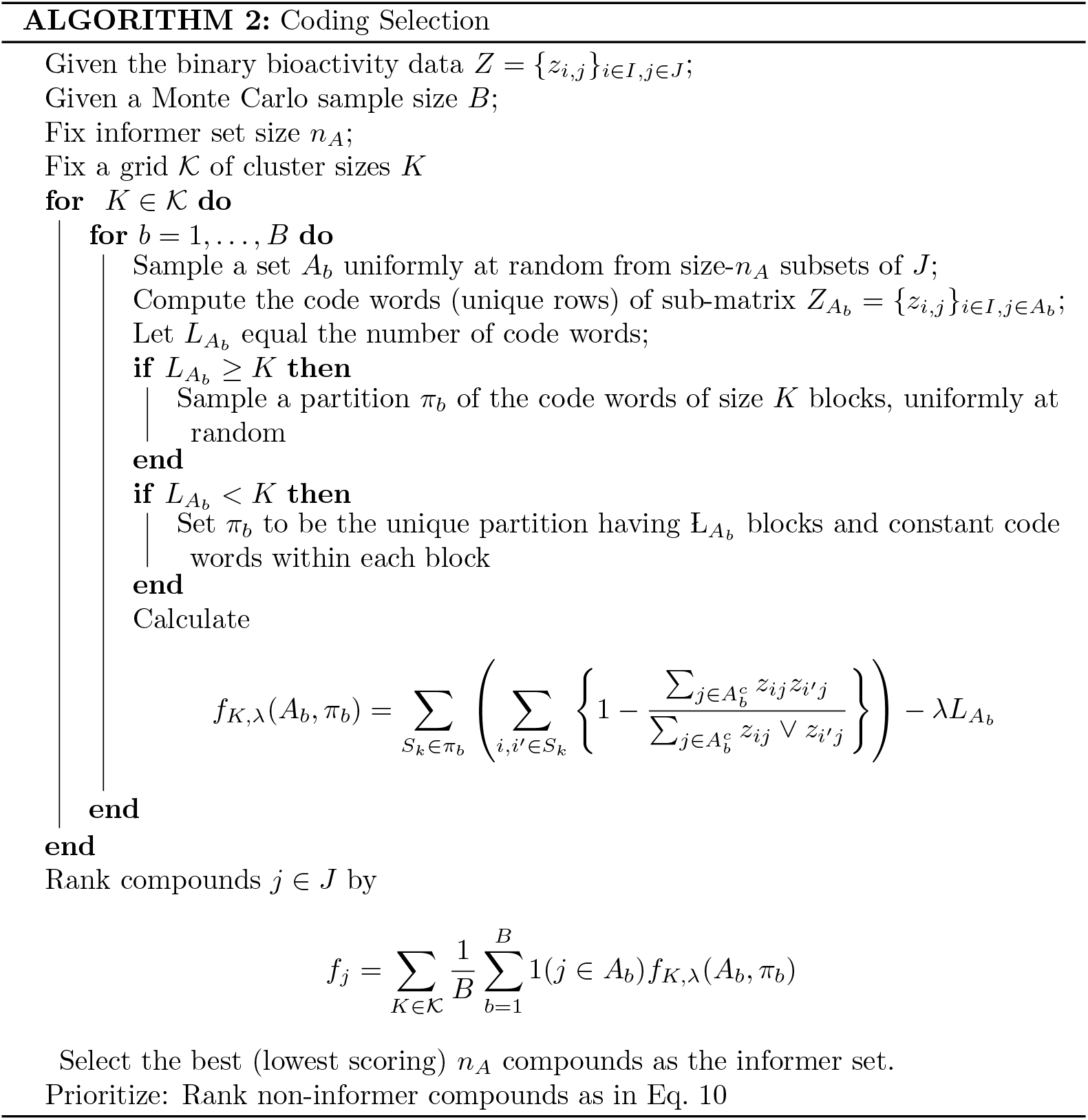

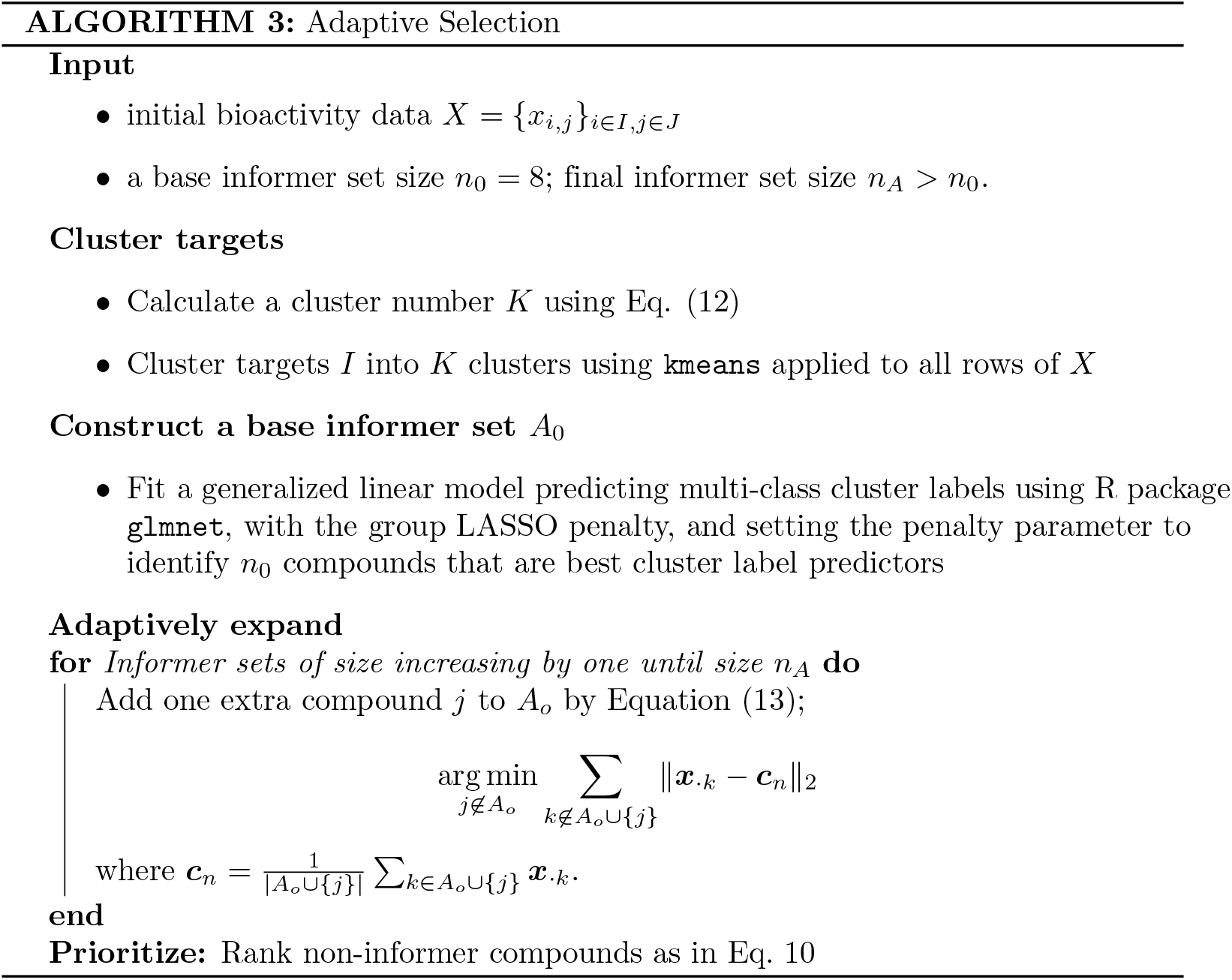

